# A novel Hox gene promoter fuels the evolution of adaptive phenotypic plasticity

**DOI:** 10.1101/2025.05.23.655234

**Authors:** Shen Tian, Bonnie Lee, Tirtha Das Banerjee, Suriya Narayanan Murugesan, Antónia Monteiro

**Affiliations:** Department of Biological Sciences, Faculty of Science, National University of Singapore; Singapore, 117543, Singapore

## Abstract

Adaptive phenotypic plasticity allows organisms to display distinct phenotypes in response to variable environments, but little is known about the genomic changes that promote the evolution of plasticity on a macroevolutionary scale. Combining spatial transcriptomics, comparative genomics, and genome editing, we show that temperature-mediated plasticity in the size of eyespot wing patterns in satyrid butterflies, a derived seasonal adaptation evolved ∼80 million years ago in this subfamily, is associated with the taxon-specific evolution of a novel promoter in the Hox gene *Antennapedia* (*Antp*). The promoter activates *Antp* expression specifically in satyrid eyespot center cells, sensitizes these cells to environmental temperatures, and plays a critical role in regulating satyrid eyespot size plasticity. Our study demonstrates that a taxon-specific cis-regulatory innovation in an ancestral developmental gene fueled the evolution of adaptive phenotypic plasticity across a clade of animals.

## Main Text

Phenotypic plasticity describes the ability of organisms to use a single genotype to produce distinct phenotypes in response to environmental cues^1,2^. When these cues are sensed during development, plasticity can alter developmental trajectories to produce fitter phenotypes adapted to each environment^3^. While we have an increasing body of knowledge how internal gene regulatory patterns change in response to external cues^4–6^, or genomic changes underlying intraspecific variations in plasticity levels on a microevolutionary scale^7–10^, we have a much fuzzier understanding of how such a complex organismal adaptation evolves on a phylogeny from ancestral, non-plastic systems. Here we use recent technical and conceptual advances in a classic model system of adaptive phenotypic plasticity – plasticity in the size of eyespot wing color pattern in satyrid butterflies – to explore the evolution of plasticity in a comparative phylogenetic framework.

The evolutionary origins and mechanisms of butterfly eyespot size plasticity has been primarily explored in *Bicyclus anynana* and its satyrid relatives. In *B. anynana*, ventral hindwing eyespot size increases with increasing environmental temperatures associated with the hot wet season (WS) and decreases with cold dry season (DS) temperatures in tropical Africa^11,12^ (Fig. 1a). This eyespot size plasticity helps individuals evade predators via different strategies in each season^13,14^, and appears to be restricted to the clade Satyrinae^15–18^, a subfamily with ∼2700 species containing *B. anynana*^19^. In *B. anynana*, hindwing eyespot size plasticity is associated with, and partially regulated by temperature-induced dynamics of an insect hormone 20-hydroxyecdysone (20E) during the larval-pupal transition^20–22^ (Fig. 1b). However, comparative work showed that variation in the titres of this hormone with temperature, and expression of the hormone receptor (EcR) in eyespot central cells, are not sufficient to explain how satyrid butterflies evolved eyespot size plasticity^17^. It is, thus, still unclear how the common ancestor of satyrids including *B. anynana* evolved temperature-mediated eyespot size plasticity.

**Fig. 1.**
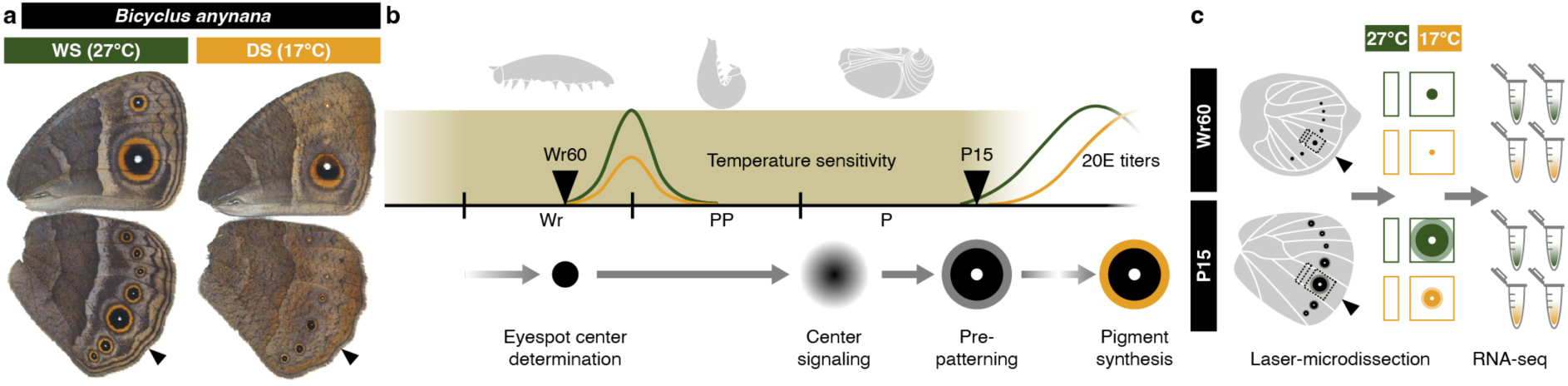
Spatial transcriptomes generated from critical developmental windows of plasticity. (**a**) The model satyrid *Bicyclus anynana* exhibits ventral hindwing eyespot size plasticity in response to rearing temperature. Arrowheads denote the Cu1 hindwing sector. (**b**) Temperature shift experiments revealed a developmental window (shaded area), when eyespot size is highly sensitive to rearing temperatures. The highly sensitive period covers essential eyespot developmental stages and temperature-induced dynamics of the insect hormone 20-hydroxyecdysone (20E), previously shown to be associated with the development of plasticity^20–22^. Within the sensitive period, two timepoints, at 60% wanderer (Wr60), and 15% pupal (P15) stages, were chosen for (**c**) the construction of spatial transcriptomes across seasonal forms, at each timepoint.

To elucidate the mechanistic evolution of eyespot size plasticity in Satyrinae, we first generated a spatial transcriptomic atlas within a refined temperature sensitive developmental window in the model satyrid *B. anynana*, aiming to resolve eyespot-specific correlates of molecular plasticity in response to temperature. From the spatial transcriptome, we identified and functionally validated a Hox gene, *Antennapedia* (*Antp*) as a conserved regulator of satyrid eyespot size plasticity. We then investigated how *Antp* regulates plasticity in a trait-and taxon-specific manner, focusing on the conservation of sequences, activities, and functions of alternative *Antp* promoters within and outside Satyrinae.

### Spatial transcriptomics from a critical time window for the development of plasticity

To resolve how external temperature cues shape internal molecular landscapes, it is essential to identify the key developmental window when eyespot size is most sensitive to rearing temperatures. Previous temperature shift experiments suggested that eyespot size is most sensitive to temperatures experienced during the final 5^th^ instar larval (L5) stage, especially the late wanderer (Wr) stage^22,23^. To confirm, as well as improve and complement these results, we repeated these experiments with a refined design (fig. S1, table S1), and found that the most temperature-sensitive developmental window is wider than previously proposed^22^, spanning Wr, pre-pupal (PP), and the first 15% pupal (P15) stages – the larval-pupal transition (Supplementary text, fig. 1b, S1). This wider period overlaps critical eyespot developmental stages: Around mid to late L5, when multiple transcription factors are expressed in the developing eyespot center cells; and the period right after pupation, when eyespot center cells signal to the surrounding cells (within 18h post-pupation, 27°C), and activate genes that prepattern the eyespot rings, and thus the final eyespot size, around P15 (∼24h post-pupation, 27°C)^24^(fig. 1b).

To construct spatial transcriptomes, we focused on one larval timepoint, 60% of the wanderer stage (Wr60), and one pupal timepoint, P15, within the temperature-sensitive window (fig. 1b). This strategy allowed us to capture both the early upstream regulators, as well as the more downstream effectors of eyespot size. Spatial transcriptomes were generated by laser microdissections of both the hindwing eyespot tissue (from the Cu1 wing sector), and an adjacent control wing tissue from the proximal side of the same sector, from both seasonal forms of *B. anynana* females, at each developmental timepoint (fig. 1c).

### Gene expression plasticity is largely systemic rather than trait-specific

To evaluate the degree of trait-specific gene expression plasticity in response to temperature, we quantified gene expression across all samples (fig. S2), and investigated if plastic genes (differentially expressed across seasonal forms, padj<0.05) in the eyespot tissue were also similarly plastic (in the same direction) in the control tissue – a systemic form of plasticity. We found that over half of the plastic genes in eyespots are systemic, whereas the rest show eyespot-specific plasticity (fig. 2a, table S2, Data S1). Gene set enrichment analyses (GSEA) were also overrepresented by the subset of genes showing systemic, rather than eyespot-specific plasticity (Supplementary text, Data S2). We hypothesized that genes with eyespot-specific plasticity or different plasticity levels were more likely to mediate the development of trait-specific phenotypic plasticity, thus we focused on those genes for subsequent investigations.

**Fig. 2.**
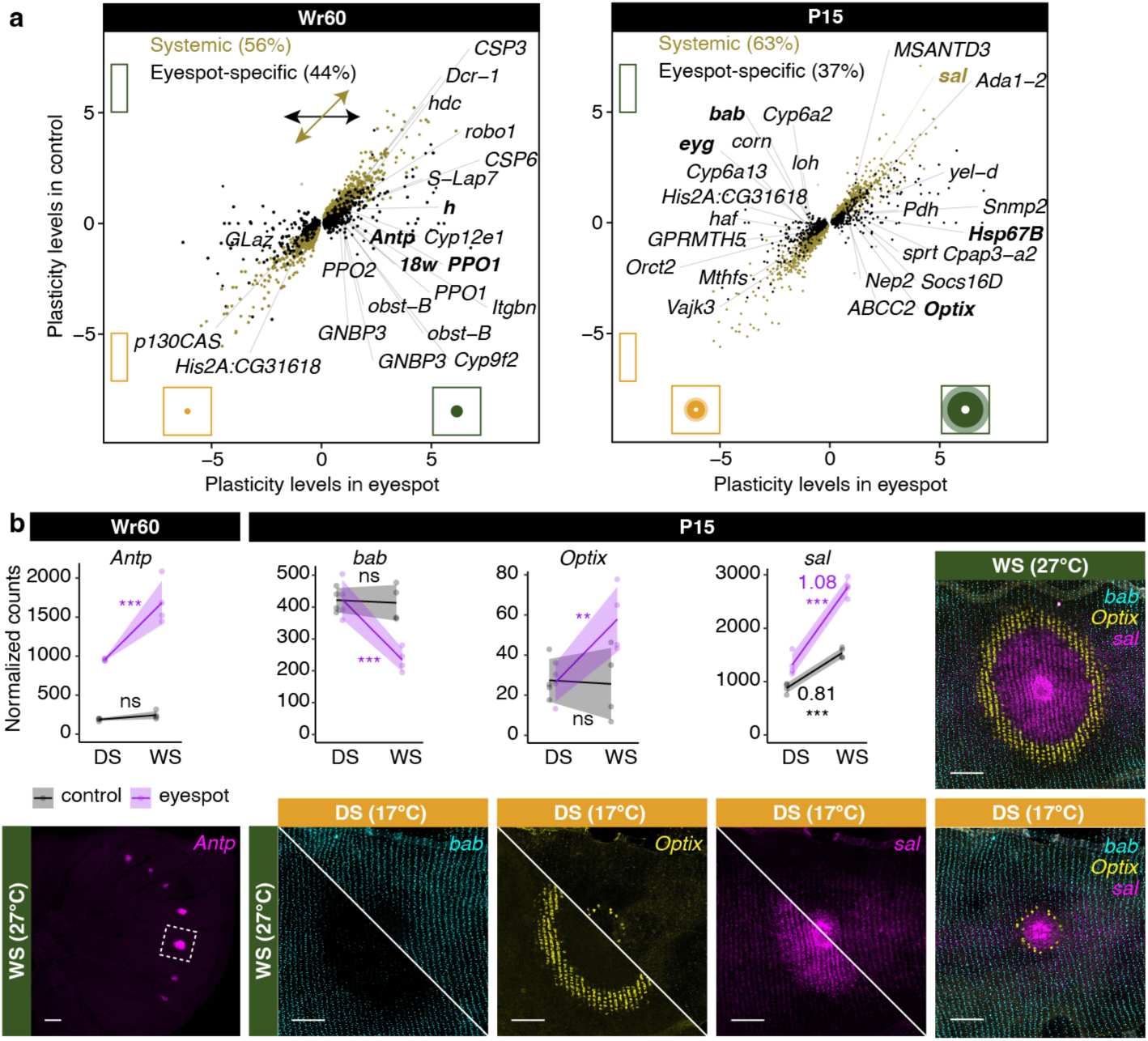
Genes showing trait-specific responses to cues are potential plasticity regulators. (**a**) Plasticity levels (log_2_ fold-changes in gene expression across seasonal forms) of the plastic genes (differentially expressed across seasonal forms, padj<0.05) in eyespots were plotted against their plasticity levels in the control tissue. Shortlisted genes with eyespot-specific plasticity were labelled (for those with known gene symbols). Candidate genes for in-vivo staining and/or functional validations are in bold. (**b**) Transcriptomic plasticity patterns of several candidate genes were well supported by HCR staining, where size of the expression domains of *bab*, *Optix*, and *sal* map to the plastic eyespot ring patterns during P15. For *sal*, different plasticity levels across tissue types are highlighted. During the early Wr60 stage, *Antp* showed eyespot-specific plasticity, with high expression levels in WS eyespot centre cells in HCR. Shaded lines: Mean values with 95CI. ns, not significant; *padj<0.05; **padj<0.01; ***padj<0.001. Scale bar: 100 microns.

### Genes showing trait-specific responses to cues are potential plasticity regulators

For ease of picking candidate genes for experimental validation, we shortlisted genes showing eyespot-specific plasticity by selecting genes with high plasticity levels (|log_2_FC|>0.8) with high significance (padj<0.01), and considerable expression levels in eyespots (mean transcript per million (TPM) >2). This generated two shortlists of candidates at each period (Wr60: 54 genes; P15: 54 genes) (fig. 2a, table S2, Data S1). From the shortlists, we picked candidates with unknown functions to visualize their spatial expression patterns using hybridization chain reaction (HCR), and to test their function using CRISPR-Cas9 mediated genome editing (Supplementary text, fig. S3-5, table S3-5, Data S3). From these experiments, we identified *bric à brac* (*bab*), a gene required to specify brown background wing color and define the outer boundary of the eyespot orange ring during P15 (fig. 2b). We also found that *Optix* and *spalt* (*sal*), two known regulators of the eyespot orange ring and black disk, respectively, exhibited eyespot-specific plasticity, with correlated size changes of the corresponding eyespot ring color elements across seasonal forms (fig. 2b). From the Wr60 list of candidates, of particular interest is *Antp,* a Hox gene essential in forewing eyespot development, but merely used to pattern the eyespot white centers and to increase eyespot size in hindwings^25^. Our transcriptome data shows that *Antp* exhibits eyespot-specific plasticity, with higher expression levels in WS, compared with DS eyespots, during Wr60 (fig. 2b). Based on its confirmed role in eyespot size regulation and its early response to temperature, we decided to further investigate the role of *Antp* in regulating eyespot size plasticity using functional tools, and in a comparative framework.

### *Antp* regulates temperature-mediated satyrid eyespot size plasticity

Previous studies mapped eyespot expression of *Antp* to the base of Satyrinae^26–28^. To investigate the role of *Antp* in regulating eyespot size plasticity, we focused on two satyrid butterflies, *B. anynana* and *Mycalesis mineus*, both increase eyespot size with increasing temperature^16^, and a non-satyrid outgroup species, *Junonia almana,* with limited decreases in eyespot size with increasing temperature^17^ (fig. 3a, f, k, S6).

**Fig. 3.**
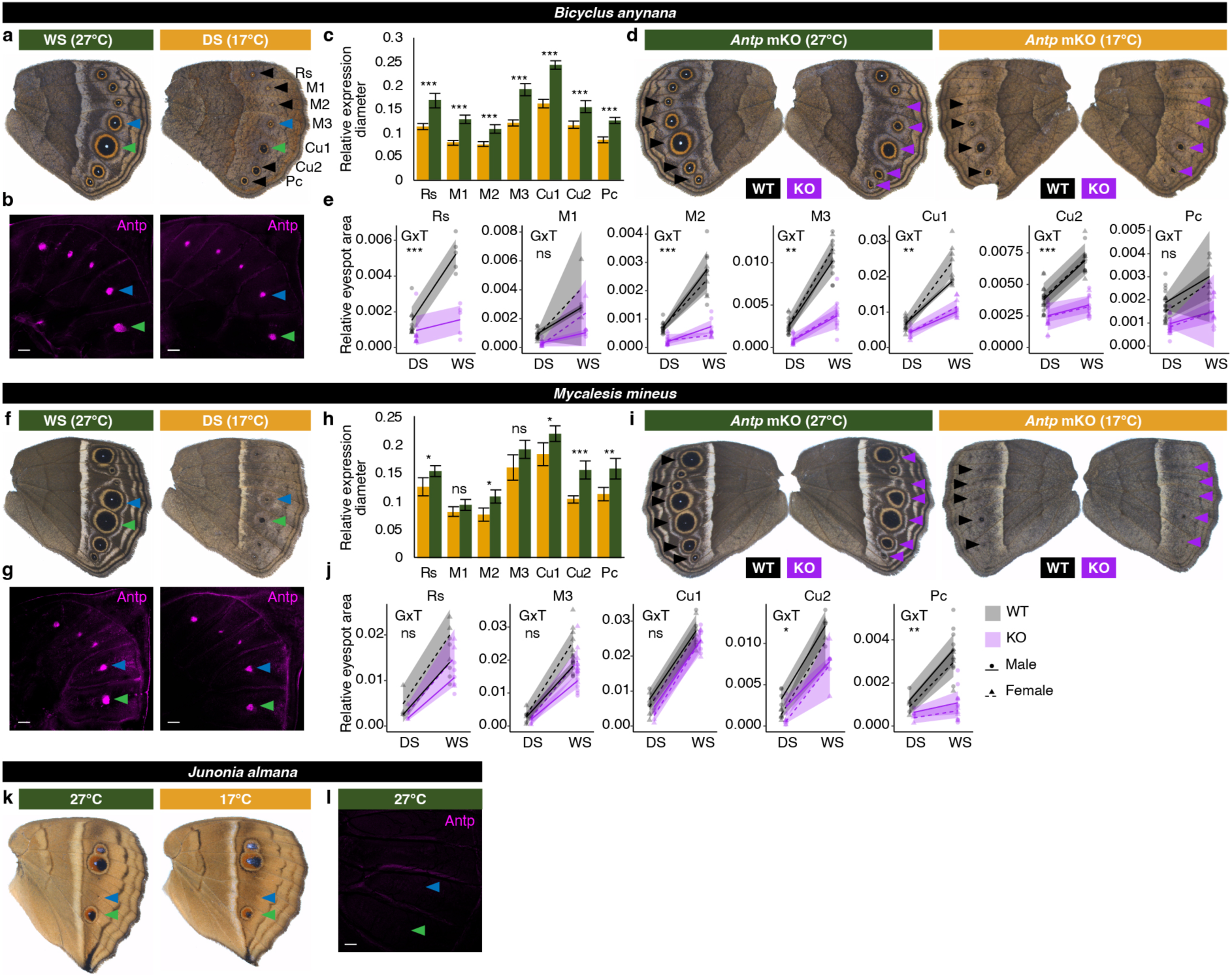
*Antp* regulates temperature-mediated eyespot size plasticity in satyrid butterflies. While satyrid butterflies, (**a-e**) *B. anynana* and (**f-j**) *M. mineus* show strong positive responses in hindwing eyespot size in response to temperature, an outgroup non-satyrid (**k-l**) *J. almana* shows limited negative responses. Immunostaining shows eyespot expression of Antp protein in (**b**) *B. anynana* and (**g**) *M. mineus* across both seasonal forms during 30-40% Wr stage, with larger expression domains in WS, compared with DS eyespot centers in (**c**) *B. anynana* and (**h**) *M. mineus*. (**l**) Eyespot expression of Antp is absent in *J. almana*. Eyespot size was quantified across mosaic knockout (mKO) *Antp* crispants of (**d**) *B. anynana* and (**i**) *M. mineus* with paired wild-type (WT) and KO eyespot, and changes in eyespot size plasticity levels was assessed in an ANCOVA, indicated by a significant (p<0.05) genotype (G) x temperature (T) interaction, in (**e**) *B. anynana* and (**j**) *M. mineus* indicates changes in levels of plasticity. Shaded line: Mean value with 95CI. Error bar: 95CI. ns, not significant; *p<0.05; **p<0.01; ***p<0.001. Scale bar: 100 microns.

We first validated the observed eyespot expression plasticity of *Antp* at the protein level, using immunostaining during the Wr stage in all three species. Antp protein was present in satyrid hindwing eyespot centers across both seasonal forms, but it was absent in the non-satyrid *J. almana* (fig. 3b, g, i, S7). At around 30-40% Wr stage, the size of the Antp protein expression domain was significantly larger in WS relative to DS eyespot centers, across all 7 eyespots in *B. anynana*, and across 5 out of 7 eyespots in *M. mineus* (fig. 3c, h, S8, table S6). This further confirmed the eyespot-specific expression plasticity of *Antp* at the protein level and suggested that this temperature-sensitive expression of *Antp* in eyespots is conserved across Satyrinae.

To test whether *Antp* regulates eyespot size plasticity, we knocked out *Antp* in both satyrids and generated mosaic knockout (mKO) crispants that we reared at 17°C and 27°C (fig. S4, table S3-4). When *Antp* was disrupted, hindwing eyespots became smaller without white centers in both species, but the size reduction was less noticeable in *M. mineus* (fig. 3d, i). Other previously reported phenotypes were also observed^25^(fig. S9-10). We quantified the hindwing eyespot area from all the mKO crispants with paired phenotypes – individual butterflies with *Antp* knock-out (KO) phenotypes on one wing, and wild-type (WT) eyespots on the other wing (fig. 3d, i) – and tested whether plasticity levels differed in WT and KO eyespots, by testing the presence/absence of a significant genotype x temperature (GxT) interaction on eyespot size. We observed significant GxT interactions for 5 out of 7 eyespots in *B. anynana*, and 2 out of 5 eyespots in *M. mineus* (only 5 eyespots are consistently visible in this species) (fig. 3e, j, table S7-8). This suggests that the recruitment of *Antp* to the eyespots of satyrid butterflies increased the size of hindwing eyespots in a temperature-dependent way, increasing their plasticity. The impact of *Antp* on the plasticity levels, however, varies across the two species examined, being stronger in *B. anynana*.

### A novel promoter *P1* activates *Antp* expression in satyrid eyespots

After confirming that *Antp* plays a conserved role in regulating eyespot size plasticity in satyrid butterflies, we sought to investigate how *Antp* gained its eyespot-specific expression to mediate trait-specific development of plasticity. For an overview of trait-specific expression patterns of *Antp*, we quantified *Antp* expression across all tissue/body parts throughout development in *B. anynana*, combining our newly generated data with a comprehensive set of published datasets^29,30^ (fig. 4a, table S9). We found that *Antp* is predominantly expressed in embryos, legs, prolegs, and wing eyespots (fig. 4b). As tissue-specific gene expression is often achieved via alternative promoters, we inspected the annotated gene model of *Antp* in the *B. anynana* ilBicAnyn1.1 genome^31^, and found nine annotated *Antp* transcripts sharing the same coding region (*CDS*), but driven by six alternative promoters, each associated with a unique non-coding first exon (fig. 4c left panel, S11). To test whether any of these alternative promoters activate *Antp* expression specifically in eyespots, we performed promoter usage analysis, to identify the proportion of total *Antp* expression driven by each promoter in each tissue (fig. 4c middle panel). We noticed that the 1^st^ distal promoter (*P1*), is exclusively activated in eyespots, and its activity can be visualised by the presence of a RNA-seq junction in eyespots that is absent in control wing tissues (fig. 4c right panel). In larval eyespots, it drives over 60% of total *Antp* expression, whereas the remaining expression is almost entirely driven by the 2^nd^ distal promoter (*P2*). In contrast to *P1, P2* is activated across all tissues/developmental stages, and drives almost the entire *Antp* expression in non-eyespot larval wing tissues (fig. 4c middle panel).

**Fig. 4.**
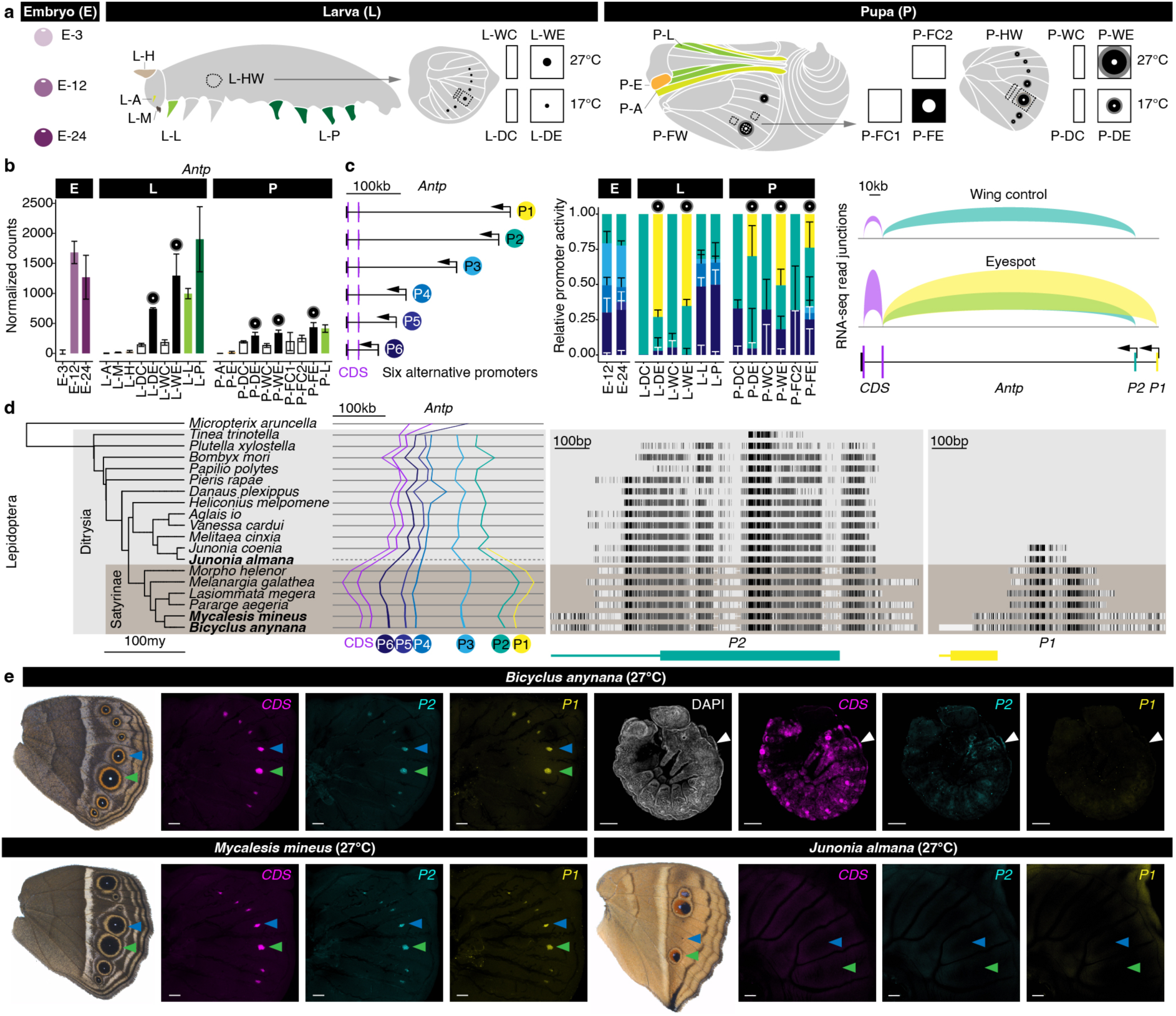
A novel promoter activates *Antp* expression in satyrid eyespots. (**a**) A transcriptomic atlas was assessed, combining our newly generated data and a comprehensive set of published tissue-specific transcriptomic data across embryonic, larval, and pupal stages. Data sources and descriptions of tissue codes are summarized in table S9. (**b**) *Antp* expression across tissue types and development. (**c**) The annotated gene model of *Antp* shows identical coding regions driven by 6 alternative promoters (left panel). The proportion of total *Antp* transcription driven by each promoter was assessed across all tissues expressing *Antp* (mid panel). RNA-seq junction reads clearly show different promoter usage across larval eyespot and non-eyespot control wing tissues (right panel). (**d**) Local synteny showing sequence conservation of all *Antp* promoters and CDSs across Lepidoptera (left panel). The highly fragmented *J. almana* genome assembly is not visualised (dotted line). Sequence conservation of the two promoters activated in larval wings, *P1* and *P2*, are highlighted (right panels). Model species used in the current study are in bold. (**e**) A direct detection of promoter activities via HCR co-staining of *Antp CDS*, *P1*, and *P2*, across *B. anynana*, *M. mineus*, *J. almana* larval wings, and *B. anynana* 48h embryos. Error bar: 95CI. Scale bar: 100 microns.

To test the extent of sequence conservation across Lepidoptera, we blasted each *B. anynana Antp* promoter and *CDS* region against genomes of all sequenced satyrids and eyespot-bearing butterflies with prior temperature-mediated plasticity data and/or eyespot expression data on *Antp*. For this purpose we also assembled and included a draft genome for *J. almana*. We found that while *Antp P2*-*6* and the two *CDS*s are all deeply conserved across Lepidoptera, *P1* is exclusively present in Satyrinae, although a fragmental orthologous sequence can also be found in the outgroup *Junonia* (fig. 4d, Data S4). This suggests that an active *P1* might have evolved exclusively in Satyrinae.

To directly detect promoter activities within and outside Satyrinae, we used HCR to co-stain the promoter-specific first exons of *P1* and *P2*, as well as *Antp CDS*, in larval wings of *B. anynana*, *M. mineus*, and *J. almana*, as well as *B. anynana* embryos (Data S3). We found that *Antp CDS*, *P1*, and *P2* were all highly activated in satyrid eyespots, and conversely, none could be visibly detected in *Junonia* eyespots (fig. 4e, S7). In *B. anynana* embryos, *Antp CDS* was present in a highly tissue/cell-specific manner across the body except the head, *P2* was activated in specific cells mainly in the T2 segment, and *P1* was not detected (fig. 4e, S12). These results suggest that a novel, taxon-specific *Antp* promoter, *P1,* actively drives *Antp* expression specifically in satyrid eyespots. Also, as *P2* is highly activated in satyrid eyespots only in the presence of an active adjacent *P1* (fig. 4e), *P1* might also serve as a novel modular enhancer, that boosts the transcriptional activity of the ancestral, pleiotropic *P2* specifically in satyrid eyespots.

### *Antp P1* regulates temperature-mediated satyrid eyespot plasticity

To examine the roles of *P1* and *P2*, we generated mKO crispants by co-injecting two guide RNAs to introduce long deletions around the conserved region of each promoter, in the model satyrid *B. anynana* (fig. 5a, S13-14, table S3-4). While *P2* mKO crispants with high-penetrance long deletions did not show visible phenotypic changes (fig. 5a right panel, S13), 27-40% *P1* mKO crispants showed smaller hindwing eyespots without white centres (fig. S14, table S4), suggesting that *P1*, but not *P2*, is necessary to mediate visibly detectable changes in eyespot size. To test if *P1* plays a role in regulating eyespot size plasticity levels, we crossed *P1* mKO crispants with WT, and then crossed two genetically identical F1 heterozygotes, with a 252bp deletion around *P1,* with each other, the resulting F2 offspring were reared across two temperatures (fig. 5a, S15, see Materials and Methods for more details). While mutant heterozygotes (*P1^+^*/*P1^Δ252^*) were visibly indistinguishable from WT (*P1^+^*/*P1^+^*) animals, mutant homozygotes (*P1^Δ252^*/*P1^Δ252^*) showed smaller hindwing eyespots without white centres (fig. 5a, S15). We compared hindwing eyespot size plasticity levels across sib-paired *P1^+^*/*P1^+^*, *P1^+^*/*P1^Δ252^*, and *P1^Δ252^*/*P1^Δ252^* animals, and found that while *P1^+^*/*P1^Δ252^* heterozygotes exhibited indistinguishable plasticity levels compared with *P1^+^*/*P1^+^*WT siblings, *P1^Δ252^*/*P1^Δ252^* mutants exhibited significantly reduced plasticity levels across 4 out of 7 hindwing eyespots compared with *P1^+^*/*P1^+^*WT siblings (fig. 5b, S16, table S10-11). This indicates that the taxon-specific *Antp P1* promotes temperature-mediated satyrid eyespot size plasticity, at least in *B. anynana*.

**Fig. 5.**
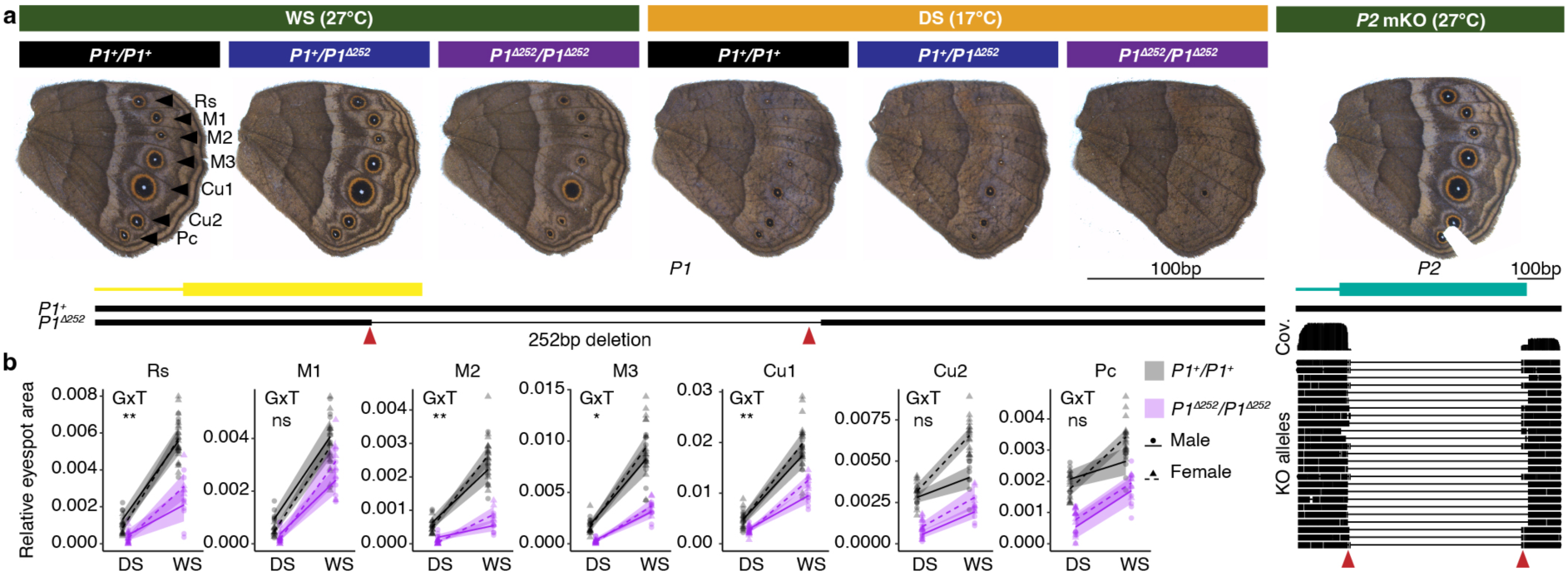
*Antp P1* regulates satyrid eyespot size plasticity. **(a)** *Antp P1* and *P2* were disrupted in *B. anynana* using CRISPR-Cas9. While mosaic knockout (mKO) crispants were generated for *P2*, a mutant line carrying a 252bp deletion was generated for *P1*. Red arrowheads denote guide RNA cut sites. *P2* mKO crispants were genotyped via Nanopore amplicon sequencing. Sequence coverage (Cov.) across the amplicon and major KO alleles are shown. **(b)** Changes in eyespot size plasticity levels were assessed across sib-paired WT (*P1^+^*/*P1^+^*) and mutant homozygotes (*P1^Δ252^*/*P1^Δ252^*) in an ANCOVA, indicated by a significant (p<0.05) genotype (G) x temperature (T) interaction. Shaded line: Mean value with 95CI. ns, not significant; *p<0.05; **p<0.01; ***p<0.001.

**Fig. 6.**
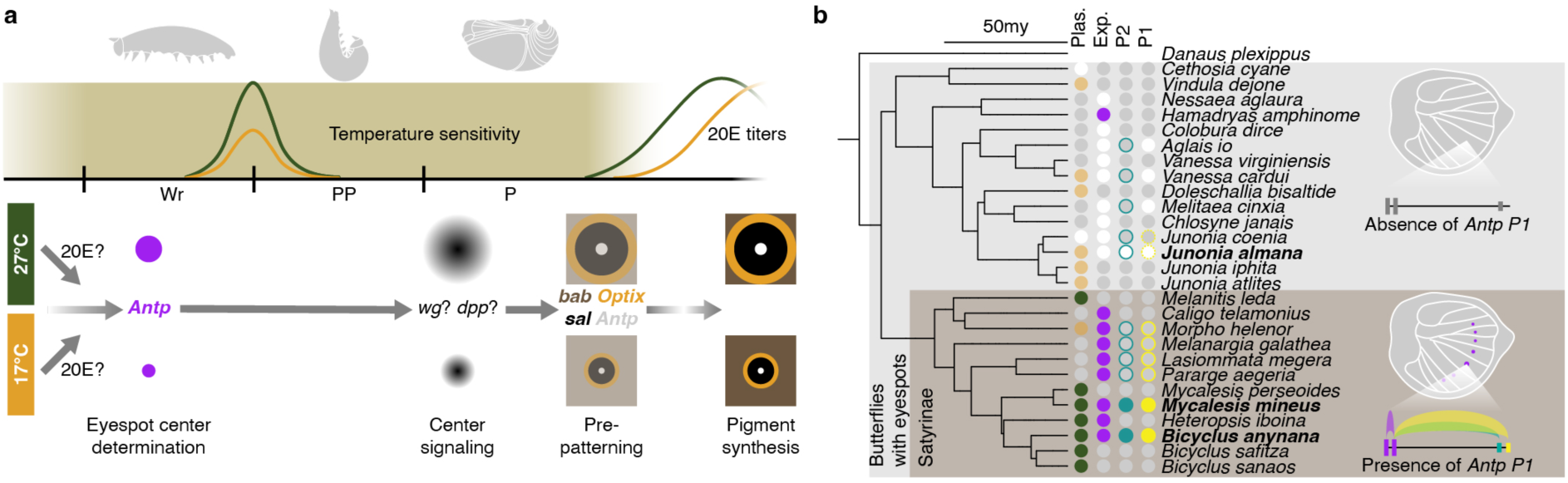
An evolutionary-developmental model of a classic system of adaptive phenotypic plasticity. We propose (**a**) a developmental model of temperature-mediated eyespot size plasticity, where *Antp* sits upstream of the eyespot gene-regulatory network (GRN), as well as an (**b**) evolutionary model illustrating how plasticity evolved on a phylogeny. The phylogeny summarizes all the current knowledge about temperature-mediated eyespot plasticity (Plas.), eyespot expression of Antp (Exp.), and sequence/activity conservation of *Antp* promoters *P1* and *P2*. The emergence of an active *P1* drives *Antp* expression in satyrid eyespots, fueling the evolution of plasticity in that clade. Plas.: Green: strong positive eyespot size response to temperature; Orange: limited negative response to temperature; White: no significant response. Exp.: Magenta: Antp protein expression in eyespots; White: no Antp eyespot expression. *P1* and *P2*: Colored outline: sequence conservation (dotted outline denotes limited conservation); Filled color: Colored: presence of promoter activities in-vivo; White: absence of promoter activities/sequence conservation. Grey: unknown.

## Discussion

In this study, we generated a spatial transcriptome to resolve trait-specific development of satyrid butterfly eyespot size plasticity. We found multiple essential regulators showing trait-specific responses to temperature across the eyespot gene regulatory network (GRN). Based on these discoveries, we propose a possible developmental model of this classic adaptive phenotypic plasticity (fig. 6a). Temperature inputs are predominantly sensed and integrated into the eyespot GRN during the larval-pupal transition, partially via altered dynamics of 20E and/or other mechanisms. This sensation and integration of cues could potentially happen at multiple timepoints within the sensitive period, and even before the surge of larval 20E titers, when *Antp* is already showing asymmetrical expression domains in eyespot center cells across seasonal forms (fig. 3). During the pupal eyespot center signaling stage, the integrated plastic responses from the upstream regulators lead to different levels of one or more signaling molecules, potentially morphogens *wingless* (*wg*) and/or *decapentaplegic* (*dpp*) from the center, that subsequently orchestrate master color regulators (*bab*, *Optix*, *sal*, *Antp*) in the surrounding cells, to prepattern the final eyespot rings with different sizes across seasonal forms (fig. 6a). Future studies are expected to elucidate the detailed molecular mechanisms.

We identified a taxon-specific *Antp* promoter that activates *Antp* expression in butterfly eyespots, which might have fueled the macroevolution of temperature-mediated eyespot size plasticity, estimated to have originated in the Satyrinae butterfly subfamily ∼80 million years ago^17^(fig. 6b). Given the polygenic nature of phenotypic plasticity^9,10^, we believe that *Antp* is not the only or major regulator of plasticity across all satyrids, evidenced by its less predominant role in *Mycalesis* with highly plastic eyespots, investigated in this study (fig. 3). Also, the neotropical *Morpho* has eyespot expression of *Antp* but shows limited temperature-mediated eyespot size plasticity^17,28^ (fig. 6b). Work in a large spectrum of satyrid species across broad geographical distributions is expected to fully understand to what extent a shared genomic innovation on seasonal adaptation could facilitate the diversification of this species-rich subfamily across heterogeneous environments.

While recent advances illustrate how nuanced modifications in ancestral cis-regulatory elements create phenotypic novelties^30,32–36^, including the adaptive circadian plasticity in *Drosophila* flies^37^, our study emphasizes the importance of novel cis-regulatory elements in fueling adaptive phenotypic evolution. Novel DNA sequences might have a higher potential to both 1) drive gene expression in novel developmental contexts^38^, and 2) bypass the pleiotropy of ancestral regulatory elements^39,40^, thereby providing a higher developmental flexibility to create phenotypic novelties. In this case, a novel cis-regulatory element is associated with the evolutionary origin of adaptive environmental sensitivity in traits.

## Acknowledgements

We thank Joseph Lee for his assistance with the promoter usage analysis. We thank Kian Long Tan for helping rear the promoter knockout lines. This work was funded by the National Research Foundation (NRF), Singapore, under its Competitive Research Program (Awards NRF-CRP20-2017-0001, and NRF-CRP25-2020-0001), and its Investigatorship program (Award NRF-NRFI05-2019-0006), and the Ministry of Education, Singapore (Award MOE-T2EP30223-0007).

## Author contributions

Conceptualization: ST, AM

Methodology: ST, TDB, AM

Investigation: ST, BL, TDB, SNM

Visualization: ST, BL

Funding acquisition: AM

Project Administration: ST, AM

Supervision: ST, AM

Writing – original draft: ST, AM

Writing – review & editing: ST, TDB, AM

## Supplementary materials

### Supplementary text

#### A refined temperature sensitive window for the development of plasticity

To precisely identify the developmental window(s) when eyespot size is most sensitive to rearing temperatures in *B. anynana*, we performed new temperature shift experiments that improved and complemented similar experiments done in the past^1,2^. An early study mapped the major temperature sensitive stage mainly from the entire 5^th^ instar larval stage (L5) to the end of 15% pupal (P15) stage^1^. However, this study did not examine any shorter yet potentially highly sensitive developmental windows within this prolonged developmental period^1^. This was addressed in a more recent study, where shorter shift windows during this period were examined, and the most temperature-sensitive stage was mapped to the end of the 5^th^ instar larval stage, the wanderer (Wr) stage^2^. However, the shift experiments were performed at a fixed time of the day (2pm) with a fixed shift duration (48h). As a result, the shift windows were not synchronized with the actual developmental transitions, which happen mostly at night in the WS form^3^. Our new shift experiments included 6 precisely staged non-overlapping shift windows synchronized with the actual developmental transitions (table S1). These shift windows covered the entire developmental trajectory from the start of L5 to the end of P15 without any gaps (fig. S1 left panel).

Females were first reared at 27°C and precisely staged for the start of the designated developmental windows, shifted to 17°C for the designated windows, or combinations of the windows, and then shifted back to 27°C (fig. S1 left panel, see Materials and Methods for more details). Adult ventral hindwing Cu1 eyespot size were measured and compared across shift schemes, where whole hindwing size was used as a covariate. The results suggested that eyespot size was sensitive to temperature from ∼50% 5^th^ instar larval stage (L5-3) to the end of P15 (fig. S1 middle panel). Within this period, three short windows, Wr, pre-pupal (PP), and P15, exhibited equally the highest sensitivity to temperature (largest relative eyespot size reduction, compared with unshifted (WS) eyespots, per unit shift time) (fig. S1 right panel). Shifting animals to low temperatures across all three stages (Wr+PP+P15, 6.8 days) produced eyespots almost the same size as DS eyespots, and indistinguishable from those that resulted from shifting the entire L5 stage (L5, 20.4 days) (fig. S1 middle panel). The latter scheme produced eyespots indistinguishable in size from DS eyespots, when the animals were reared entirely at 17°C (fig. S1 middle panel).

### Gene expression plasticity in a plastic trait is largely systemic rather than trait-specific

We examined the extent that plastic genes in eyespots also showed plastic responses (in the same direction) in control tissues, at each of the two developmental stages. We found that with the exception of a few genes (less than 10 in each stage) showing plasticity in opposite directions across tissue types, more than half of the plastic genes in eyespots (wanderer stage: 56%; early pupal stage: 63%) were also plastic in control tissues, representing systemic plasticity across the wing. The rest (wanderer stage: 44%; early pupal stage: 37%) were plastic only in eyespots (fig. 2a, table S2, Data S1). To elucidate whether the two gene categories also diverge in gene functions, we performed gene set enrichment analysis (GSEA) on the entire set of plastic genes in eyespots, the subset showing systemic plasticity, and the subset showing eyespot-specific plasticity (Data S2). We found that more than half of the GO terms enriched (padj<0.1) in the entire set of plastic genes were shared by terms enriched in the subset showing systemic plasticity (wanderer stage: 64%, 202 of 314; early pupal stage: 59%, 178 of 302), while only a smaller proportion was shared by those enriched in the subset showing eyespot-specific plasticity (wanderer stage: 35%, 110 of 314; early pupal stage: 31%, 95 of 302). The two subsets showing systemic and eyespot-specific plasticity were enriched with largely distinct GO terms, with very few terms in common (wanderer stage: systemic: 246, eyespot-specific: 152, common: 31; early pupal stage: systemic: 259, eyespot-specific: 141, common: 19). This suggests that molecular plasticity in gene expression is largely systemic, not only at the level of individual genes, but also at the level of gene functions, in the plastic eyespot tissues.

### Genes with trait-specific expression plasticity are potential plasticity regulators

From the shortlist of genes showing eyespot-specific plasticity at the Wr60 stage, we picked four candidates. In addition to the Hox gene *Antennapedia* (*Antp*), with known expression patterns and roles in eyespot development^4^, three were not investigated before: a sensory bristle pre-pattern gene *hairy* (*h*), a Toll-like receptor*18-wheeler* (*18w*), and an enzyme involved in would healing *prophenoloxidase 1* (*PPO1*) (fig. 2a left panel, fig. S3). We investigated the spatial expression of these three genes via HCR in WS larval wings, at both Wr60 and an earlier mid L5 stage. Their HCR staining appeared irrelevant to eyespot development (fig. S3 upper and mid panels, Data S3). Knocking them out in WS animals either produced no visible phenotypes (*18w*, *PPO1*), or phenotypes irrelevant to eyespots (*h*) (fig. S4-5, table S3-5).

From the P15 stage, we picked three genes that were not investigated before: a *Heatshock protein 67B* (*Hsp67B*), a BTB transcription factor *bric à brac* (*bab*), and a Pax-homeobox transcription factor *eyegone* (*eyg*) (fig. 2a right panel, fig S3). We investigated their expression patterns via HCR in both seasonal forms at P15 stage (fig. S3 lower panel, Data S3). While HCR staining of *Hsp67B* and *bab* aligned well with the transcriptomic data, no visible signal was detected for *eyg* (fig. 2b, S3 lower panel). *Hsp67B* maps to the eyespot center, with larger expression domain in WS compared with DS eyespot centers (fig. S3 lower panel). Knocking it out in WS animals produced no visible phenotypes (fig. S4, table S3-4). *Bab* maps to the background brown wing color outside the eyespot golden ring, with a larger *bab*-negative eyespot area in the WS compared with DS animals, corelated with the final eyespot size. (fig. 2b). We also knocked out *eyg* in WS animals and surprisingly, we observed comet-like eyespots with eyespot foci elongated to the wing margin, and in some cases, eyespots became bigger (fig. S4-5, table S3-5). As this is a role more relevant to eyespot foci determination during L5, we stained *eyg* also in WS larval wings, at both mid L5 and Wr60, and noticed that it is highly expressed in the wing margin, but its expression levels drop towards the developing eyespot foci (fig. S3 upper and mid panels). This suggests that *eyg* represses eyespot focal signaling derived from wing margin early in development, although our transcriptomic data suggested that this gene does not show any expression plasticity during Wr60 (fig. S3 mid panel).

## Materials and Methods

### Insect husbandry

Lab populations of *B. anynana* butterflies have been reared in the laboratory since 1988, originally collected in Malawi. *M. mineus* butterflies were collected from Clementi Forest, Singapore, *J. almana* butterflies were collected from Seletar West Farmway, Singapore, both under a National Parks Board permit (NP/RP14-063-7a).

All the butterflies were reared in two climate rooms, at 17°C and 27°C, leading to the development of DS form and WS form, respectively. Both climate rooms have a 12:12 day: night cycle (daytime and nighttime start from 7am and 19pm, respectively), with 60% relative humidity. *B. anynana* and *M. mineus* larvae were fed young corn leaves and adults were fed mashed banana. *J. almana* larvae were fed *Ruellia repens* and adults were fed artificial nectar.

### Temperature shift experiments

Temperature shift schemes included individual or combinations of 6 non-overlapping short developmental windows spanning the entire 5th instar larval stage and the first 15% pupal stage in *B. anynana* (fig. S1). Females were reared at 27°C, shifted to 17°C for the designated developmental windows, or combinations of windows, followed by a shift back to 27°C.

For the schemes involving wanderer, pre-pupal, and pupal stages, the start and end timepoints of these stages were precisely scored for each individual using the time-lapse function of an Olympus Tough TG-5 camera^3^. One exception was shifting the entire 5th instar larval stage (L5). The pupation timepoints of this scheme were largely dispersed among individuals due to the long rearing durations (∼20 days) at 17°C. For this scheme, individual pupation status was checked manually every 6 hours throughout the day and fresh pupae were shifted back to 27°C in a timely basis.

For the schemes involving pre-wanderer 5th instar larval stages (L5-1, –2, and –3), 4th larval ecdysis could happen throughout the day, so 4th instar larvae were checked every 6 hours throughout the day. If they entered 5th instar larval stage between 0 am-12 pm of a day, that day was considered day 0 of the 5th instar larval stage, if they entered 5th instar larval stage between 12 pm-24 pm, the next day was considered day 0. The shift start time of these schemes was 0 am at midnight, and shift durations lasted 5 days. Since the developmental pace of the pre-wanderer stages could vary by 1-2 days among individuals, animals with the actual shift schemes overlapping more than 50% of the adjacent shift schemes were excluded as they did not represent a genuine effect of the designated shift schemes anymore.

The temperature shift experiments were performed by grouping 3-8 females perfectly synchronized in development in small plastic containers. Each shift scheme included at least 20 females, except for L5-3, where 12 females were used. Statistics of the temperature shift experiments were summarized in table S1.

Adult butterflies were frozen around 4h after adult eclosion. Wings were dissected and imaged with a Leica DMS 1000 microscope, using 1.25 x magnification level and a Leica 0.32 x Achromat objective lens. Ventral hindwing Cu1 eyespot areas and whole hindwing areas were measured in pixels using the Quick Selection Tool in Photoshop.

### Calculation of temperature sensitivity in eyespot size

As different temperature shift schemes involved different shift durations, we define temperature sensitivity in eyespot size as reductions in relative eyespot size, compared with unshifted (WS) eyespots, per unit shift time (h). Under this definition, relative eyespot size (eyespot area divided by hindwing area) was calculated for Cu1 eyespots across the 6 non-overlapping shift schemes (L5-1, –2, –3, Wr, PP, P15), as well as in the WS form. Temperature sensitivities in eyespot size were then calculated as:

Temperature sensitivity = (Mean relative eyespot size of WS – relative eyespot size of the shift scheme) / Mean shift duration

Temperature sensitivity values were then compared across the 6 non-overlapping shift schemes in a statistical test.

### Laser microdissection and RNA sequencing

Seasonal forms of *B. anynana* were precisely staged as previously described^3^. Female hindwings were dissected from WS and DS forms at 60% wanderer (Wr60) and 15% pupal (P15) stages. Laser microdissection was performed according to a published protocol^5^. In brief, freshly dissected wings were immediately mounted on PEN membrane slides on ice, then fixed and dehydrated using ice-cold ethanol and acetone, and stained using ThermoFisher Arcturus HistoGene Staining Solution. Fixed and dehydrated tissue slides were kept at –80 °C till enough samples were collected. Laser microdissection was performed to cut the eyespot tissue and an adjacent non-eyespot control tissue from the hindwing sector Cu1. For the Wr60 stage, we cut a rectangular area within the Cu1 sector for the eyespot tissue, centered on Cu1 eyespot center. For the P15 stage, we cut an inscribed circular area within Cu1 sector to sample the entire eyespot pattern, centered on the Cu1 eyespot center. In both developmental stages, we cut a narrow rectangular area just next to the proximal side of the Cu1 eyespot region within the Cu1 sector, as a control wing tissue. One biological replicate included microsections pooled from 30 hindwings from 15 individuals (both sides) for the Wr60 stage, or 8 hindwings from 8 individuals (one side only) for the P15 stage. Four biological replicates were included for each condition. For a brief tissue digestion prior to RNA extraction, microsections were collected and pooled in lysis buffer, and digested using proteinase K at 55 °C for 9min. Total RNAs were extracted using mirVana miRNA Isolation Kit following the manufacturer’s instructions.

Quality check for total RNA samples was performed using Nanodrop 2000, gel electrophoresis, and LabChip. Poly-A enriched stranded mRNA sequencing libraries were constructed using VAHTS Universal V8 RNA-seq Library Prep Kit. For Illumina sequencing, over 30 million 150 bp read pairs per sample were generated using Illumina NovaSeq platform. Quality check, library construction, and sequencing were carried out by Azenta, Singapore.

### Gene expression and functional enrichment analysis

Trimmomatic 0.39 was used to trim adaptors from raw sequencing reads (options: PE ILLUMINACLIP:TruSeq3-PE-2.fa:2:30:10:8:true MAXINFO:40:0.2 MINLEN:32). Adaptor-trimmed clean reads were used to quantify gene expressions (NCBI *B. anynana* v1.2 genome) using Salmon 1.6.0, with the quasi-mapping mode (options: –-validateMappings –-seqBias – gcBias). Gene expression levels were normalized, and differential gene expression analysis was performed using DESeq2 in R studio. Numbers and lists of differentially expressed genes (padj<0.05) in multiple comparisons were summarized in table S2 and Data S1.

Gene set enrichment analysis (GSEA) was performed using Babelomics 5 for the entire list of plastic genes (differentially expressed across seasonal forms, padj<0.05) in eyespots, and for the two subsets of the list showing systematic and eyespot-specific gene expression plasticity, at each developmental stage. Lists of enriched gene ontology (GO) terms (padj<0.1) were summarized in Data S2.

### Promoter usage analysis

Promoter usage analysis was performed across the newly generated data and a comprehensive set of published tissue/body part-specific transcriptomic data^6,7^ (table S9). Adapters were trimmed from raw sequencing reads as described above. Adaptor-trimmed clean reads were used to quantify gene expressions (ilBicAnyn1.1 genome^8^) using Salmon 1.6.0, with the quasi-mapping mode (options: –-validateMappings –-seqBias –gcBias). Gene expression levels were normalized using DESeq2 and the expression levels of *Antp* were checked across all samples.

To quantify splice junctions for the estimation of promoter activities, adaptor-trimmed clean reads were mapped to the ilBicAnyn1.1 genome using STAR 2.7.10a. A two-pass approach was adopted to index the genome for a more sensitive junction discovery, as described in the STAR manual. In brief, the ilBicAnyn1.1 genome was first indexed with the genome annotations for the 1^st^ mapping pass across all the samples with usual STAR parameters (options: –-outSJfilterReads Unique). The splice junctions detected from the 1^st^ pass were merged and filtered. All the non-canonical junctions, and junctions supported by too few reads (reads<=2) were discarded. The consolidated and filtered junctions were used as annotated junctions to index the genome again with the genome annotation, and the new genome index was used for a 2^nd^ mapping pass (options: –-outSJfilterReads Unique –-outFilterMultimapNmax 1). Junction files generated from the 2^nd^ mapping pass were used as inputs for promoter usage analysis using proActiv^9^ in R studio. Relative promoter activities – the proportion of total *Antp* transcription initiated by each promoter, were quantified across all samples with sufficient *Antp* expression levels and junction reads to support the analysis.

### *J. almana* genome sequencing and assembly

Genomic DNA was extracted from thorax of a female *J. almana* adult using Omega E.Z.N.A Tissue DNA Kit. Library preparation and sequencing for Illumina short-read paired-end (2 × 150 bp) sequencing were performed by Genewiz, China, resulting in 20 GB of raw data.

Raw FASTQ files were trimmed and quality-controlled using the bbduk script from BBMap tools. Short-read-based genome assembly was performed using Platanus 1.2.4 with default settings. The genome was first assembled into contigs using the platanus assemble command, followed by scaffolding of the assembled contigs. Finally, gaps were closed using the platanus gap_close command, resulting in a final assembly size of 476 MB.

### In-situ HCR

Detection of candidate genes and promoters was carried out based on the protocol described^10^ with few modifications. For protein-coding genes, probes were designed on the coding sequences (CDS). For *Antp* promoters, probes were designed on an extended genomic region around the unique 1^st^ exon initiated by each promoter. All the HCR probe sets were listed in Data S3. Briefly, wings or embryos were dissected in 1x PBS and transferred to glass chambers where they were fixed in 1x PBST supplemented with 4% formaldehyde for 30 mins. After fixation, the tissues were treated with a detergent solution^11^ and washed first with 1x PBST followed by washes in 5x SSCT. Afterward, embryos were embedded in polyacrylamide gel^12^ and transferred to a confocal dish, while wings were transferred to glass wells with 500µL of 30% probe hybridization buffer. The hybridization reaction involved incubation in a solution containing 50μL (50μM) of probe set against each gene (IDT) in 2000 µL of 30% probe hybridization buffer followed by rigorous washing with 30% probe wash buffer. Afterward, tissues were washed with 5X SSCT and incubated in amplification buffer for 30 mins. For the chain reaction, a solution with HCR hairpins (Molecular instruments) in amplification buffer was added to the tissues followed by washes in 5x SSCT. Finally, the tissues were mounted on an inhouse mounting media and imaged under an Olympus fv3000 confocal microscope. The primary incubation was carried out for 16-24h at 32 °C, and the secondary hairpin incubation for 8h at room temperature.

### Immunostaining

*B. anynana* and *M. mineus* female hindwings were dissected at ∼30-40% wanderer stage from both seasonal forms, *J. almana* female hindwings were dissected at the same stage, from WS form only. Fresh wings were fixed with 4% formaldehyde in cold fix buffer (0.1M PIPES pH 6.9, 1mM EGTA pH 6.9, 1% Triton x-100, 2mM MgSO4) for 30min. Fixed wings were washed four times with cold PBS and incubated in block buffer (50mM Tris pH 6.8, 150mM NaCl, 0.5% IGEPAL, 5mg/mL BSA) overnight at 4 °C. Wings were then incubated with primary antibody diluted in wash buffer (50mM Tris pH 6.8, 150mM NaCl, 0.5% IGEPAL, 1mg/mL BSA) for 24h at 4 °C. A mouse anti-Antp 4C3 antibody (Developmental Studies Hybridoma Bank, final dilution 1:200) was used as the primary antibody. For *J. almana*, an additional guinea pig anti-Sal GP66.1 antibody^13^ (final dilution 1:20000) was used as a positive control. Wings were washed 4 times for 20min each with cold wash buffer and then incubated with secondary antibody diluted in wash buffer for 2h at 4 °C in a dark environment. For *B. anynana* and *M. mineus*, Alexa Fluor 555 Goat anti-Mouse was used as the secondary antibody (final dilution 1:500). For *J. almana*, Alexa Fluor 488 Goat anti-Mouse and Alexa Fluor 555 Goat anti-Guinea Pig (final dilution 1:500) were used for a double staining of Antp and Sal. Wings were washed 4 times for 20min each with cold wash buffer, mounted on glass slides, and kept at 4 °C before imaging. Ventral surfaces of the wings were imaged using an Olympus FV3000 confocal laser scanning microscope with identical configurations.

### Calculation of eyespot expression diameter of Antp protein

For each hindwing eyespot of *B. anynana* and *M. mineus*, two orthogonal diameters of the Antp protein expression domain were measured, one in parallel with the proximal-distal axis of the sector, and the other perpendicular to it (fig. S8). The size of the Antp protein expression domain, defined as eyespot expression diameter, was calculated as the mean of the two measured diameters. The distance between the Cu1 and M3 eyespot centers was also measured and used as a hindwing size proxy, which was entered in the statistical analysis as a covariate. All diameters and distances were measured in pixels using the ruler tool in Photoshop. To avoid bias, all images were measured ‘blindly’, without knowledge of the treatment group they belonged to.

### CRISPR-Cas9 genome editing

Embryonic CRISPR-Cas9 knock out (KO) experiments were performed in *B. anynana* and *M. mineus* following the established protocol^14^. Guide RNAs were designed using CRISPRdirect. Specificity of the guide RNA target sequences was checked by blasting against the NCBI *B. anynana* v1.2 genome or the published *M. mineus* genome^15^. For protein-coding genes, template single guide DNAs (sgDNAs) were produced by PCR, using NEB Q5 High-Fidelity DNA Polymerase. sgDNAs were transcribed into sgRNAs using NEB T7 RNA Polymerase. sgRNAs were purified by ethanol precipitation. Size and integrity of sgDNAs and sgRNAs were checked using gel electrophoresis. For *Antp* promoters, synthetic crRNAs were directly ordered from Integrated DNA Technologies (IDT). Target sequences of all guide RNAs were listed in table S3.

For microinjection of guide RNAs, corn leaves were placed in *B. anynana* or *M. mineus* adult cages around 2-3pm. Leaves were left in the cages for 1h and eggs were collected and placed onto 1mm-wide double-sided tapes in petri dishes. For protein-coding genes, we used a final concentration of 500ng/μL IDT Alt-R S.p. Cas9 Nuclease V3 (with 1 x Cas9 buffer), and 250ng/μL sgRNA in the injection mixture. The injection mixture was incubated at 37 °C for 10 min prior to injection. 0.5 μL food dye was added per 10 μL mixture to facilitate the visualization of the injected solutions. Different sgRNAs for the same protein-coding gene were not mixed but injected independently. For *Antp* promoters, eggs were injected with CRISPR/Cas9 protein-crRNA-tracrRNA complexes. Stock solutions of crRNAs and tracrRNA were prepared as 1000ng/μL, 4μL of crRNA were incubated with 4μL of tracrRNA and 12μL of Nuclease-Free Duplex Buffer at 95°C for 5 mins and then left to cool down at room temperature. We used a final concentration of 500ng/μL Cas9 protein (with 1 x Cas9 buffer), and ∼ 200ng/μL annealed crRNA-tracrRNA in the injection mixture. Two crRNAs designed for each promoter were co-injected to introduce long deletions around the promoter regions.

Microinjection was performed using glass capillary needles. After injection, moistened cotton balls were placed into each Petri dish. Hatchlings were shifted to corn plants to complete development. Adults were frozen ∼6h after adult emergence. Phenotypic changes were inspected manually and mutants were imaged using a Leica DMS 1000 microscope. The CRISPR injection statistics were summarized in table S4.

### Quantification of eyespot size from mosaic *Antp* mutants with paired phenotypes

CRISPR-Cas9 mediated *Antp* KO experiments were performed in *B. anynana* and *M. mineus*. Injected eggs were reared at either 27°C or 17°C. Adult butterflies were collected and manually screened for all visible phenotypes. To examine the effect of *Antp* KO on the eyespot size plasticity levels, all crispants with paired WT and KO hindwing eyespot phenotypes – individual butterflies with *Antp* KO eyespot phenotypes on one wing, and WT phenotypes on the other, were used. Wings were imaged as described above. To get precise measurements of all eyespots, especially for the smaller and DS eyespots, wings were imaged with the same configurations, with a 4 x magnification level for zoom-in views of eyespots, as well as a 1.25 x level for whole hindwings. Eyespot areas of each ventral hindwing eyespot with paired WT and KO phenotypes, as well whole hindwing areas, were measured in pixels in Photoshop, as described above. Eyespot size was measured across all 7 hindwing eyespots in *B. anynana* and 5 out of 7 hindwing eyespots in *M. mineus*, as *M. mineus* M1 and M2 eyespots and/or their associated white eyespot centers were not consistently observed even among WT animals. Number of crispants with paired phenotypes was summarized in table S7.

### T7 endonuclease assay

T7 endonuclease assays were performed to confirm the in-vivo cutting efficiencies of all sgRNAs before large-scale egg injections. Genomic DNA was extracted from ∼20 injected eggs, 4 days after injection, using E.Z.N.A Tissue DNA Kit. For each target gene, one pair of genotyping (GT) primers was designed to amplify a 300∼600bp genomic region spanning the cut sites using PCRBIO Taq Mix Red. PCR products were purified using ThermoFisher GeneJET PCR Purification Kit. 200ng of PCR products were denatured and re-hybridized with NEB Buffer 2 in a total volume of 20μL, following the reported temperature settings^14^. The product was divided into two tubes, one treated with 1μL T7 endonuclease (New England Biolabs), and the other without endonuclease as a control. Both were incubated at 37 °C for 15min and were subsequently run on a 2% agarose gel. All guide RNAs showed efficient cutting of the designated target sites in-vivo, except one *Hsp67B* guide in *B. anynana* (fig. S4). GT primers were listed in table S3.

### Detect long deletions by PCR

For *Antp P1* and *P2*, efficient in-vivo genomic editing by the co-injected crRNAs and the resulting long deletions were confirmed by PCR before large-scale egg injections. Genomic DNA was extracted from injected eggs using E.Z.N.A Tissue DNA Kit. GT primers (table S3) were designed to amplify a ∼400-700bp genomic region flanking both crRNAs for each promoter using PCRBIO Taq Mix Red. PCR products were checked using gel electrophoresis for the presence of long deletion alleles (fig. S13-14).

### Genotyping *eyg* and *h* mosaic mutants via Sanger sequencing

On-target mutations across mosaic knock out (mKO) mutants of *eyg* and *h* were checked using Sanger sequencing. Genomic DNA was extracted from a single wing showing mutant phenotypes from each mKO mutant using E.Z.N.A Tissue DNA Kit. The same GT primers used in the T7 assay were used to amplify the genomic region spanning the target sites using PCRBIO Taq Mix Red. PCR products were purified using ThermoFisher GeneJET PCR Purification Kit, and were sent for Sanger sequencing by 1^st^ Base (Axil Scientific Pte Ltd, Singapore). Guide-induced indels were checked using the Synthego ICE Analysis tool v3. Genotyped individuals and their indel rates were shown in table S5.

### Genotyping *Antp P2* mosaic mutants via Nanopore sequencing

Since *Antp P2* mKO mutants did not exhibit any visible phenotypic changes, on-target long deletions were first checked with PCR. Genomic DNA was extracted from right hindwings of eight mKO females using E.Z.N.A Tissue DNA Kit. GT primers used for genotyping injected eggs were used to amplify the genomic region flanking the two crRNAs using NEB Q5 High-Fidelity DNA Polymerase, and PCR products were purified using ThermoFisher GeneJET PCR Purification Kit. PCR products were checked using gel electrophoresis and NanoDrop, and three samples showing clear long deletions (fig. S13) were sent for Nanopore amplicon sequencing, to generate 1Gb data for each sample. Quality check, library preparation, and sequencing were carried out by Azenta, Singapore. Nanopore long reads were mapped to the reference sequence of *Antp P2* from the ilBicAnyn1.1 genome using minimap2 2.26, and alignments were visualized in Integrated Genomics Viewer (IGV) (fig.5, S13).

### Generation of *Antp P1* mutant line and eyespot quantification

Both crRNAs targeting *Antp P1* were injected to generate F0 *P1* mKO mutants. F0 mKO females showing eyespot phenotypes were crossed with WT males. Some of these F0 females might carry one or several mutant alleles in the germline and passed the mutant alleles to F1. The F1 offspring were screened via hemolymph extraction and Sanger genotyping, as described below.

To screen F1, around 10μL of hemolymph was extracted from 5th instar larvae and suspended in 200μL Saline-Sodium Citrate buffer. Hemocytes were collected by centrifugation, then resuspended and incubated in 20μL digestion buffer (1.1mL 1M Tris-HCl of pH6.8 in 50mL deionized water) with 2μL proteinase K (New England Biolabs) at 37 °C for 15min. Upon heat inactivation of proteinase K at 95 °C for 3min, 3μL of the cell lysate containing genomic DNA was used for PCR using designed GT primers (table S3). PCR products were run on the gel, F1 heterozygotes carrying long deletions showed two bands, the upper WT bands and the lower mutant bands. To isolate different long deletion alleles, the mutant bands were excised from the gel, purified using ThermoFisher GeneJET Gel Extraction Kit, and sent for Sanger Sequencing by 1st Base (Axil Scientific Pte Ltd, Singapore). Among all mutant alleles, we obtained at least two F1 heterozygotes of the opposite sex carrying a frequently occurring 252bp deletion.

The two F1 heterozygotes were crossed with each other, and the resulting F2 eggs were equally split and reared at either 27°C or 17°C, generating sib-paired mutant homozygotes, heterozygotes, and WT, in both seasonal forms. F2 adults were frozen ∼6h after adult emergence for 1) genotyping and 2) eyespot quantification. Heads were used for DNA extraction, PCR, and gel electrophoresis for genotyping purposes, as described above. A sample gel image showing bands of mutant homozygotes, heterozygotes, and WT is shown in fig. S15.

Hindwings were imaged as described above, with a 4x magnification level for zoom-in views of eyespots, as well as a 1.25 x level for whole hindwings. Eyespot areas of each ventral hindwing eyespot, as well whole hindwing areas, were measured in pixels in Photoshop, as described above. Eyespots were measured ‘blindly’, without knowledge of the genotype they belonged to. The number of F2 animals analyzed and their genotypes were summarized in table S10.

### Sequence conservation of the *Antp* features

Sequence conservation of *Antp CDSs* and promoters (*P1*-*6*) were checked across Lepidoptera, including all sequenced satyrids and other eyespot-bearing butterflies with published data on temperature-mediated eyespot size plasticity^16,17^, and/or eyespot expression of *Antp*^13,18,19^. *B. anynana* ilBicAnyn1.1 genomic sequences of *Antp* features were blasted against the selected lepidopteran genomes to check for the presence of any orthologous sequences in each genome. Species, genome versions, and coordinates of the blasted hits were summarized in Data S4.

### Statistical analysis

A one-way analysis of covariance (ANCOVA) was used to assess the impact of temperature shift schemes on 1) the size of the ventral hindwing Cu1 eyespot 2) the temperature sensitivity of the ventral hindwing Cu1 eyespot size, and 3) the impact of rearing temperatures on the eyespot expression diameter of Antp protein in each ventral hindwing eyespot. In the case of 1) and 3), hindwing area and hindwing size proxy, respectively, was used as a covariate. Homogeneity of variances was first tested using Levene’s test. Based on the test results, data was square root transformed to reduce skewness, when necessary. A one-way ANCOVA was performed in R studio with type III sums of squares. For 1) and 2), Tukey’s multiple comparison test was used to detect which shift schemes produced significant (p<0.05) changes in eyespot size or temperature sensitivity.

A two-way ANCOVA was used to assess the impact of genotype (G), temperature (T), and their interactions (GxT) on the eyespot size from 1) *Antp* mKO crispants and 2) sib-paired *Antp P1* KO lines, controlled for hindwing area and sex as covariates. Homogeneity of variances was first tested using Levene’s test and the data was square root transformed, when necessary. A two-way ANCOVA was performed in R studio with type III sums of squares, using the model:

Eyespot_area ∼ Wing_area + Sex + Temperature*Genotype

For the analysis mentioned above, figures were made using means and 95% CIs of the ratio of eyespot area (or eyespot expression diameter) / wing area (or wing size proxy) to visualize the distribution of the original datapoints. Statistical test results were shown in fig.3,5, S1, S16, and summarized in table S6, S8, S11.

### Supplementary Figures

**fig. S1.**
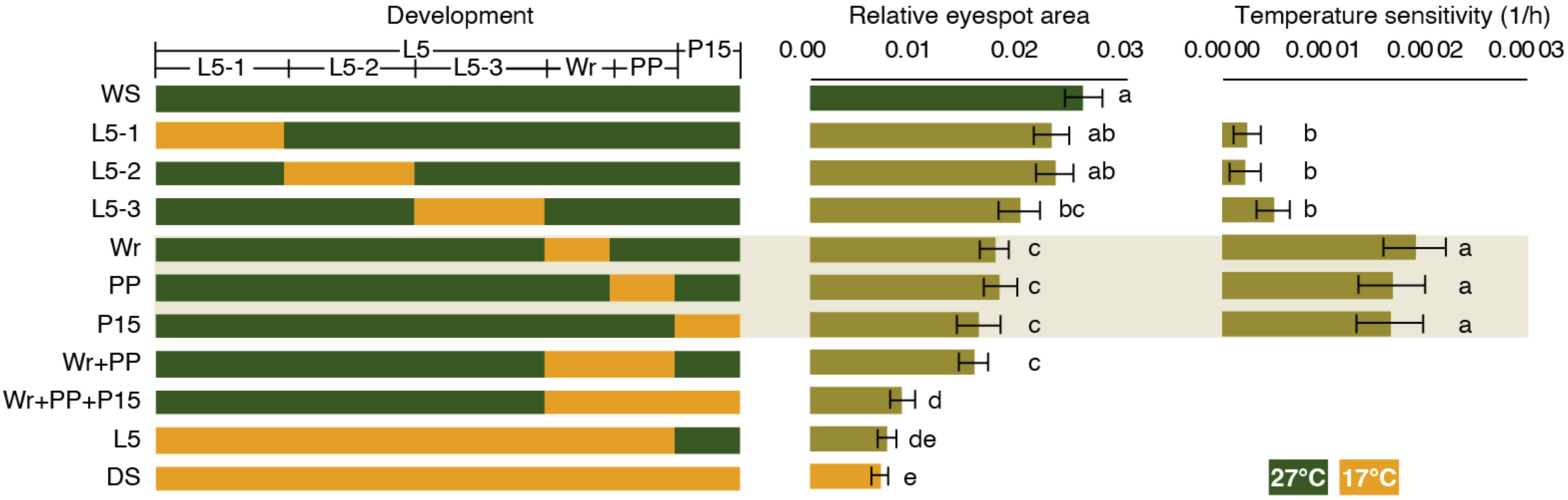
Temperature shift experiments identified a prolonged temperature sensitivity window. Shift schemes in refined temperature shift experiments are illustrated (left panel). Ventral Cu1 eyespot areas (middle panel), and temperature sensitivities (right panel, only across the six non-overlapping windows) were quantified and compared across the schemes. Shift schemes with the same letter are not significantly different from each other, as determined by Tukey’s test. Error bar: 95CI. WS, wet season, DS, dry season; L5, 5^th^ instar larva; Wr, wanderer, PP, prepupa; P, pupa.

**fig. S2.**
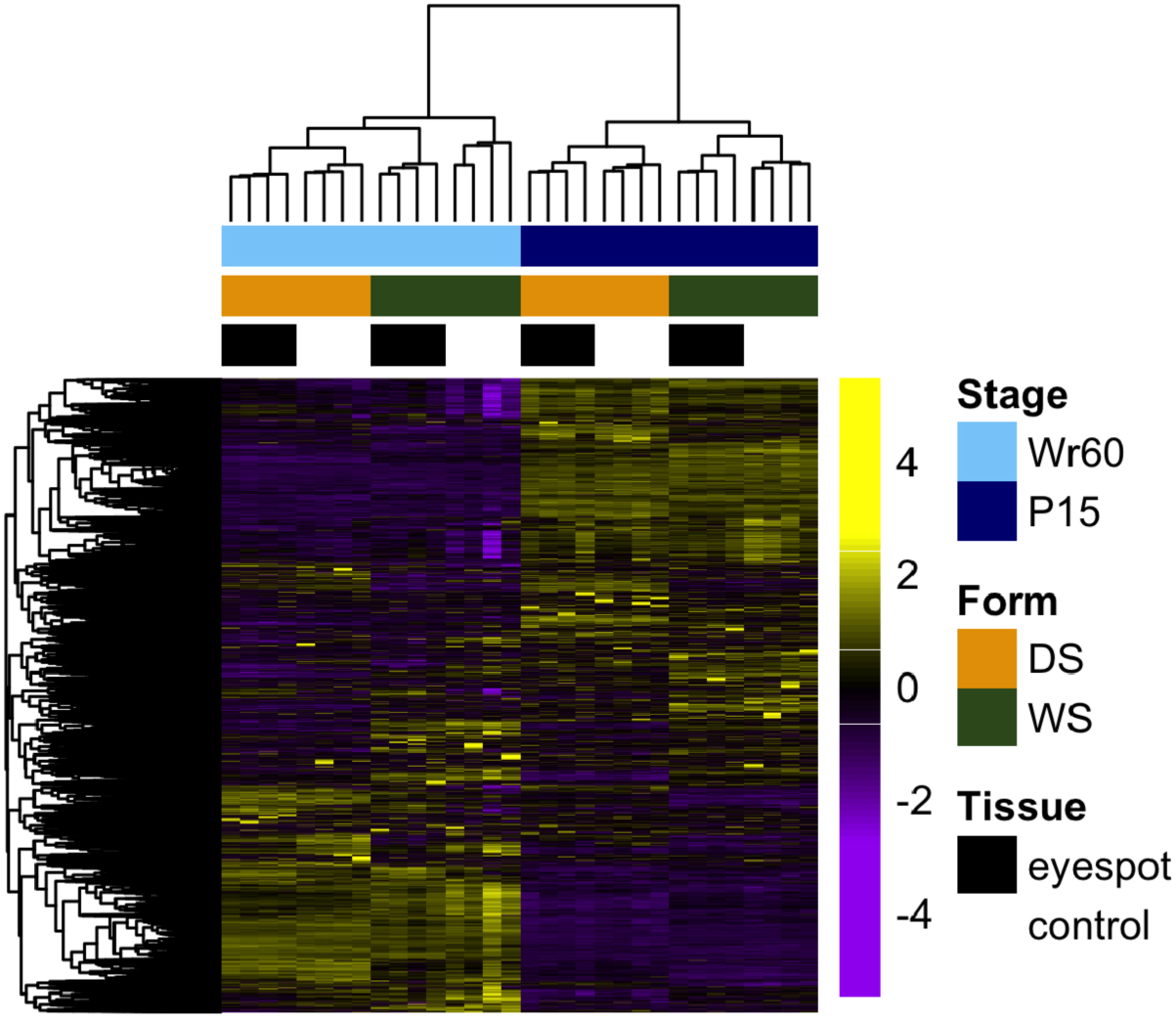
Hierarchical clustering of the laser-dissected spatial transcriptome. The heatmap shows clean clustering of the biological replicates primarily by developmental stages, then by seasonal forms, and tissue types.

**fig. S3.**
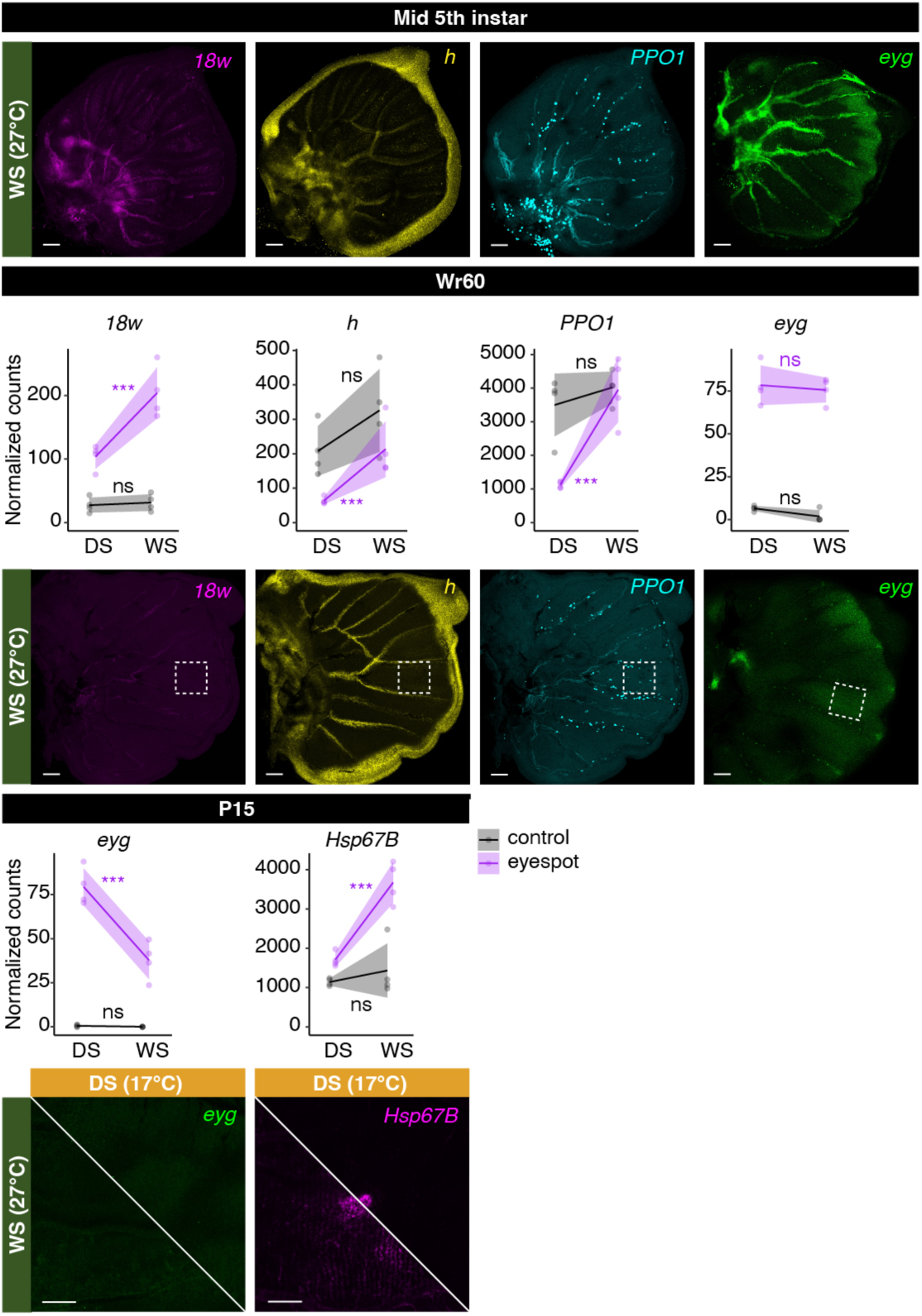
Expression patterns of the candidate plasticity regulators. Candidate genes were picked from the spatial transcriptomes as the data suggests that these genes exhibit eyespot-specific expression plasticity. HCR staining was performed to validate their spatial expression patterns, highlighting the eyespot regions. Shaded lines: Mean values with 95CI. ns, not significant; *padj<0.05; **padj<0.01; ***padj<0.001. Scale bar: 100 microns.

**fig. S4.**
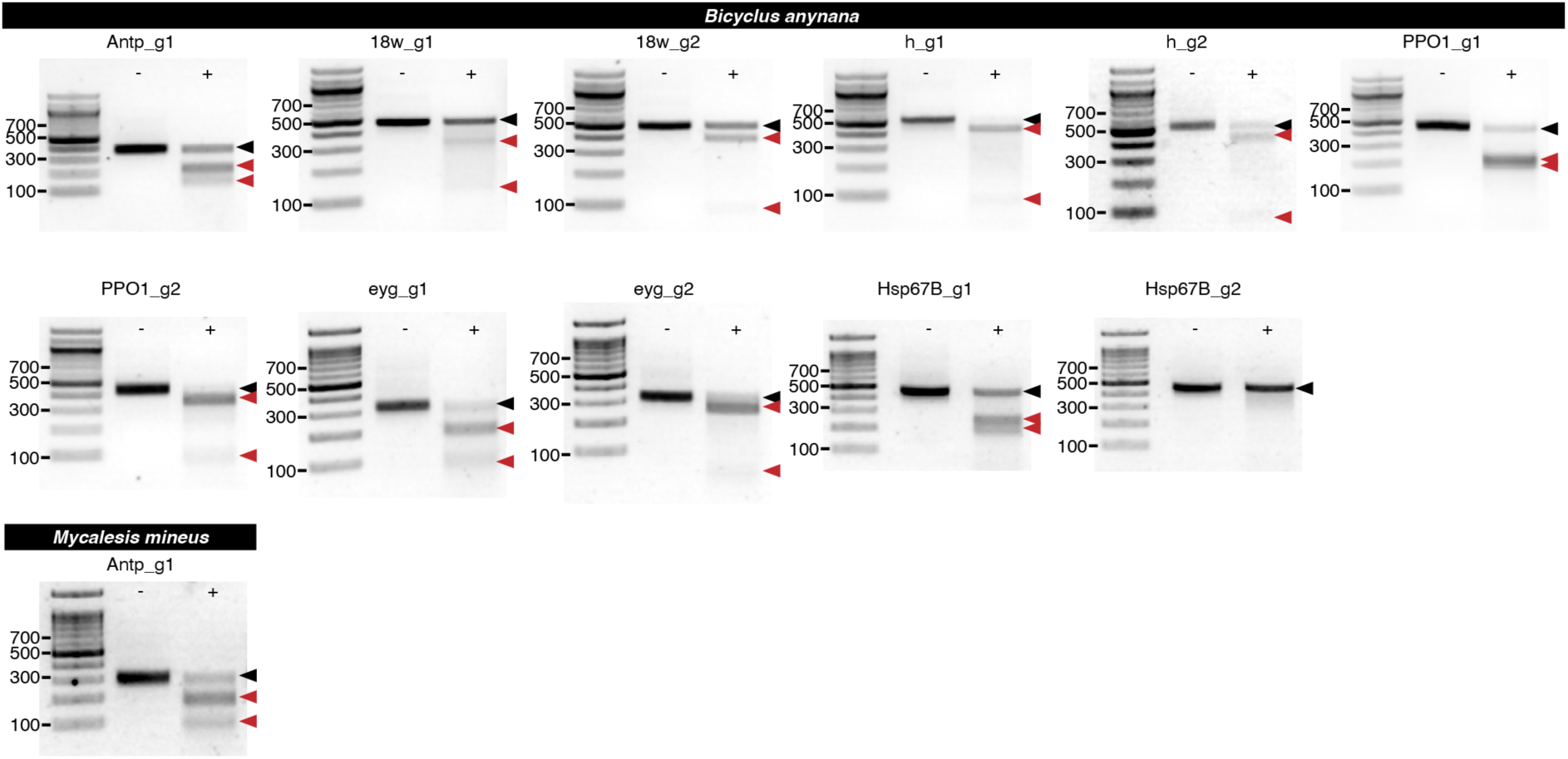
T7 endonuclease assay. A T7 endonuclease assay was performed to test the cutting efficiencies of all sgRNAs in-vivo. Control wells without T7 endonuclease treatments are denoted ‘-’, while treatment wells are denoted ‘+’. The uncut bands is denoted by a black arrowhead, while the two fragmental bands after T7 endonuclease digestion are indicated by red arrowheads. Among all guides, only Hsp67B_g2 did not show successful cutting.

**fig. S5.**
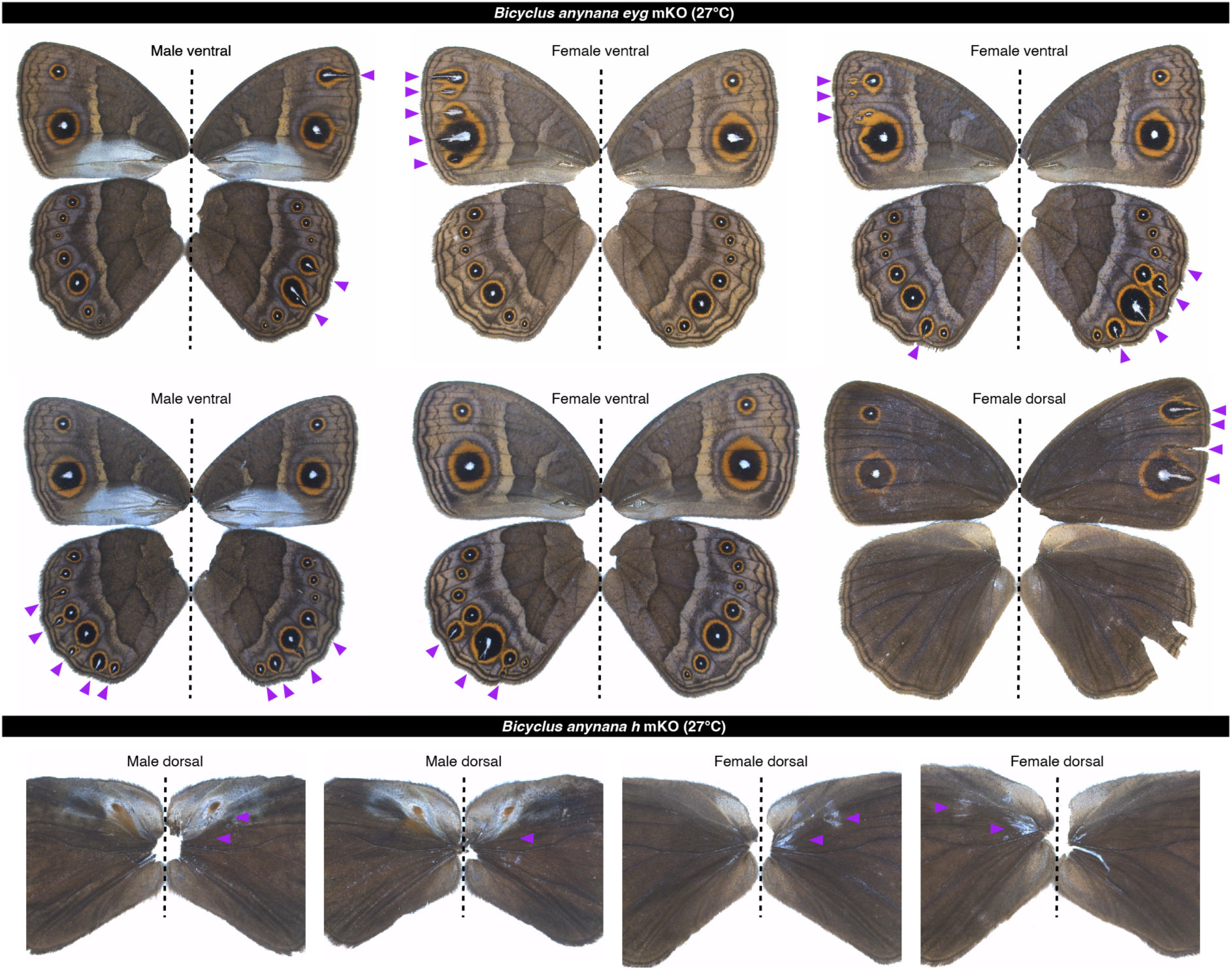
Representative phenotypes of *eyg* and *h* mKO crispants in *B. anynana*. A dotted line separates left and right sides of the same individual. Purple arrowheads denote mutant phenotypes.

**fig. S6.**
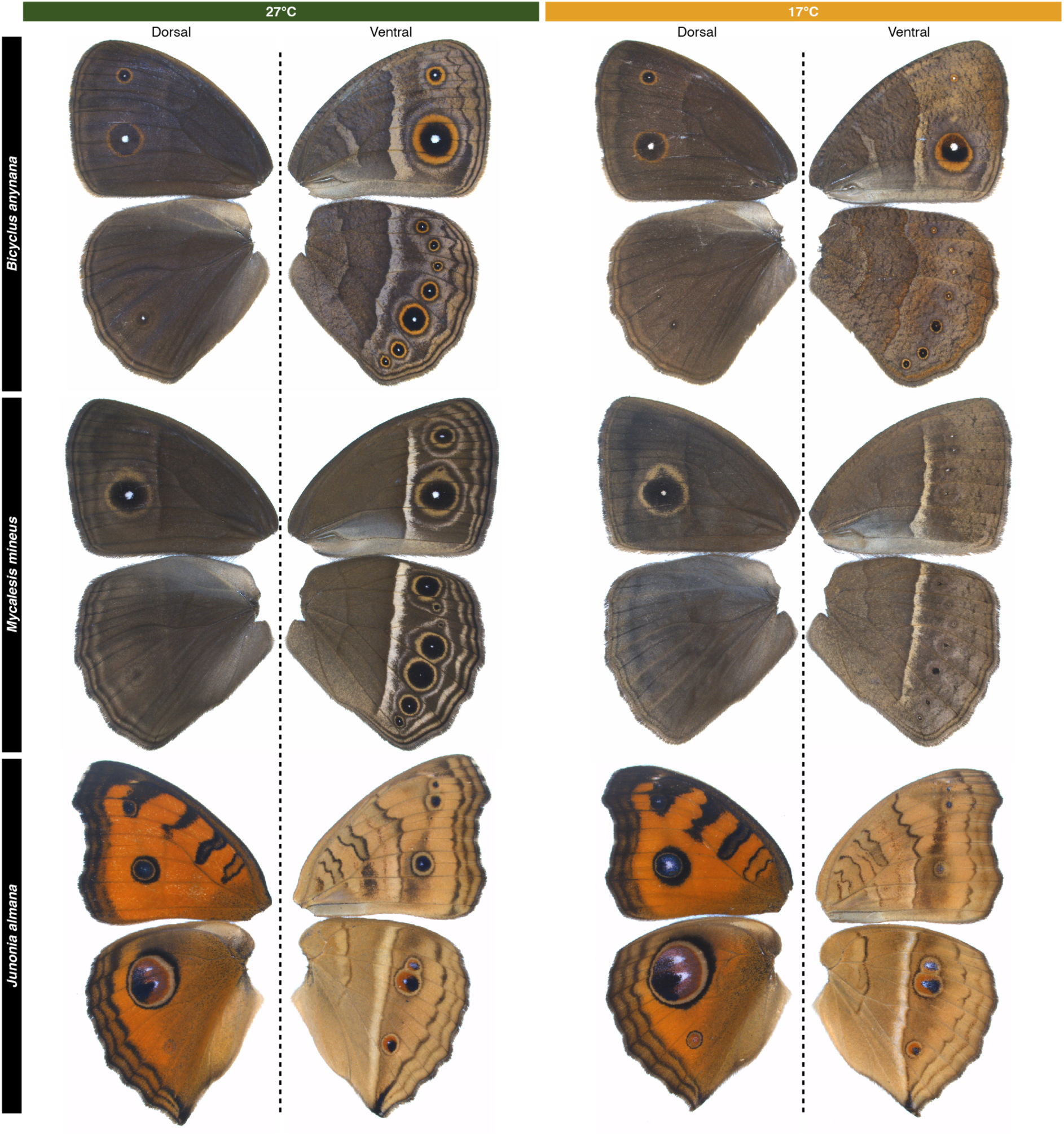
Representative phenotypes of the three model butterflies reared at two temperatures. A dotted line separates dorsal (left) and ventral (right) sides of the same female individual.

**fig. S7.**
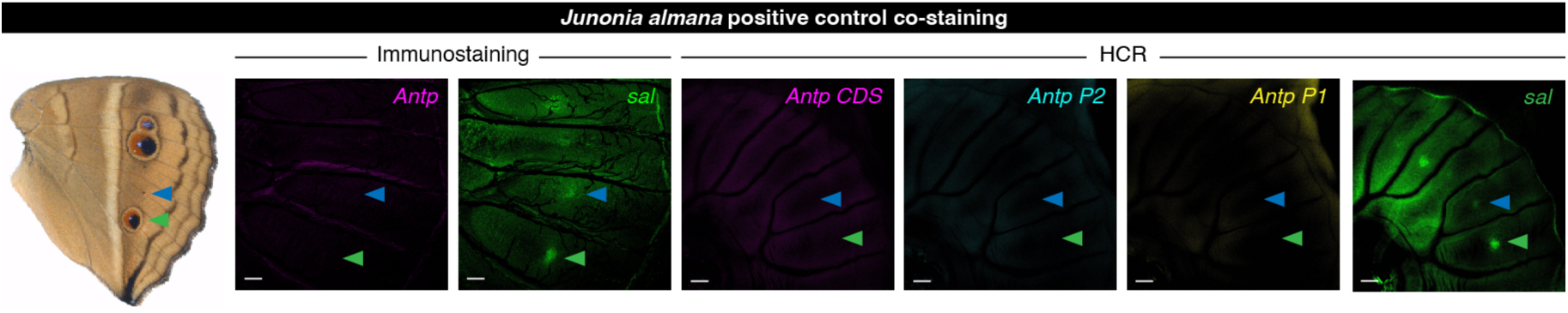
Co-staining of a positive eyespot marker in *J. almana*. *Sal* was used as a positive eyespot marker in both immunostaining and HCR.

**fig. S8.**
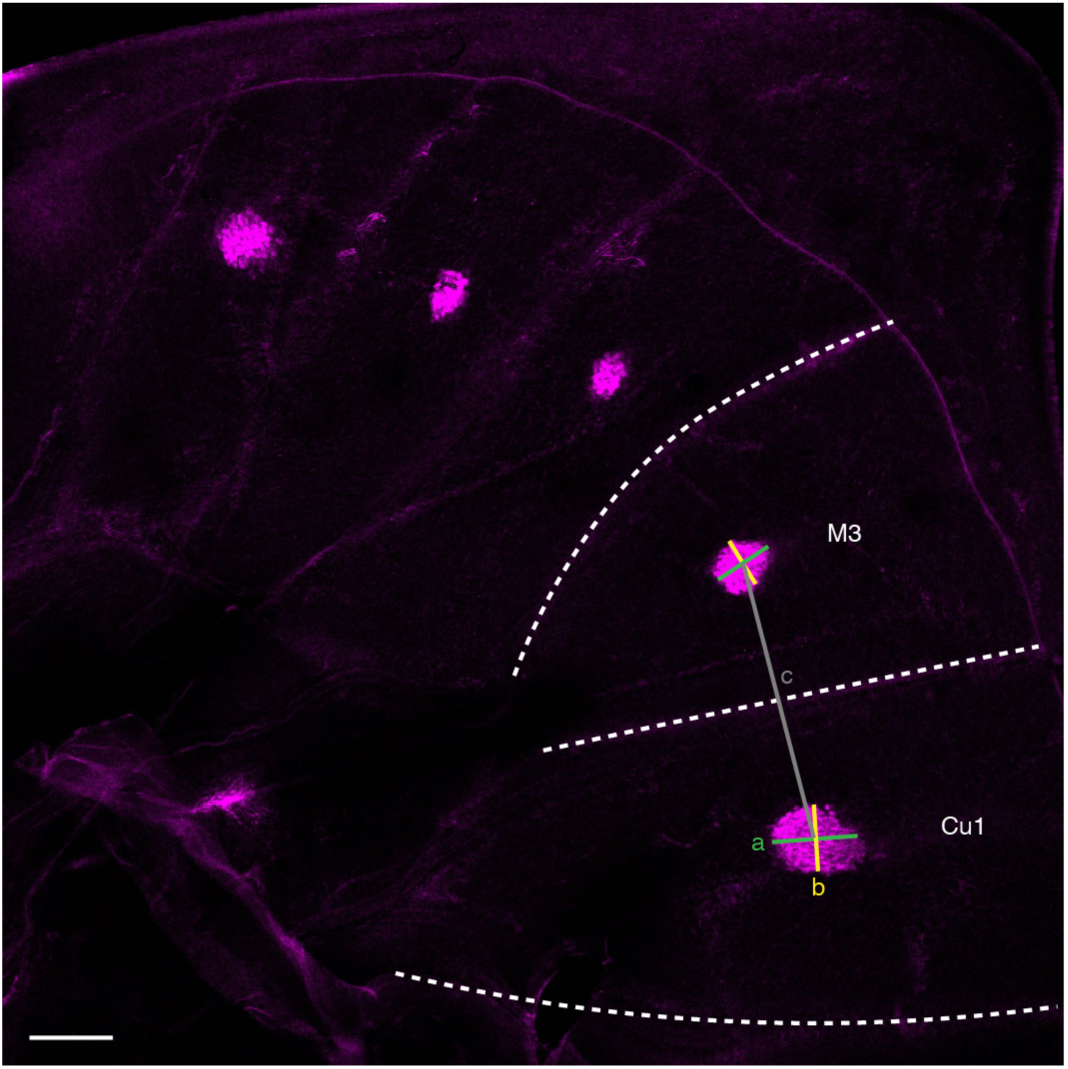
Calculation of Antp protein eyespot expression diameter. Size of the Antp protein expression domain in eyespots was calculated as eyespot expression diameter. For each ventral hindwing eyespot (here we use Cu1 eyespot as an example), eyespot expression diameter was calculated as the mean of the two diameters of the Antp protein expression domain (a and b), one (a) was parallel to the proximal-distal axis of the sector (outlined in dotted lines), while the other (b) perpendicular to it. The distance between M3 and Cu1 eyespot centers (c) was constantly used as a hindwing size proxy, and entered as a covariate in statistical analyses. Scale bar: 100 microns.

**fig. S9.**
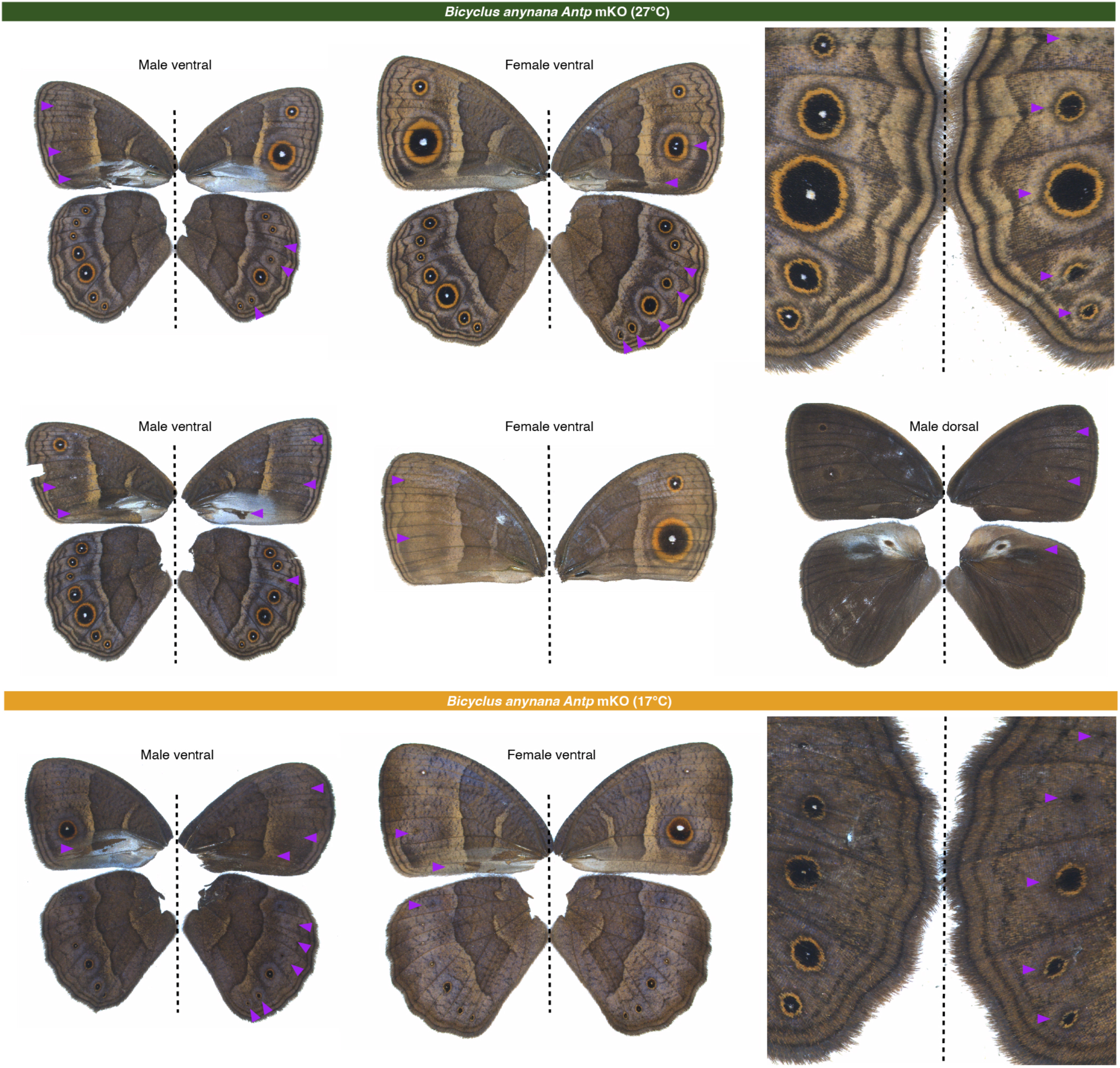
Representative phenotypes of *Antp* mKO crispants in *B. anynana*. A dotted line separates left and right sides of the same individual. Purple arrowheads denote mutant phenotypes.

**fig. S10.**
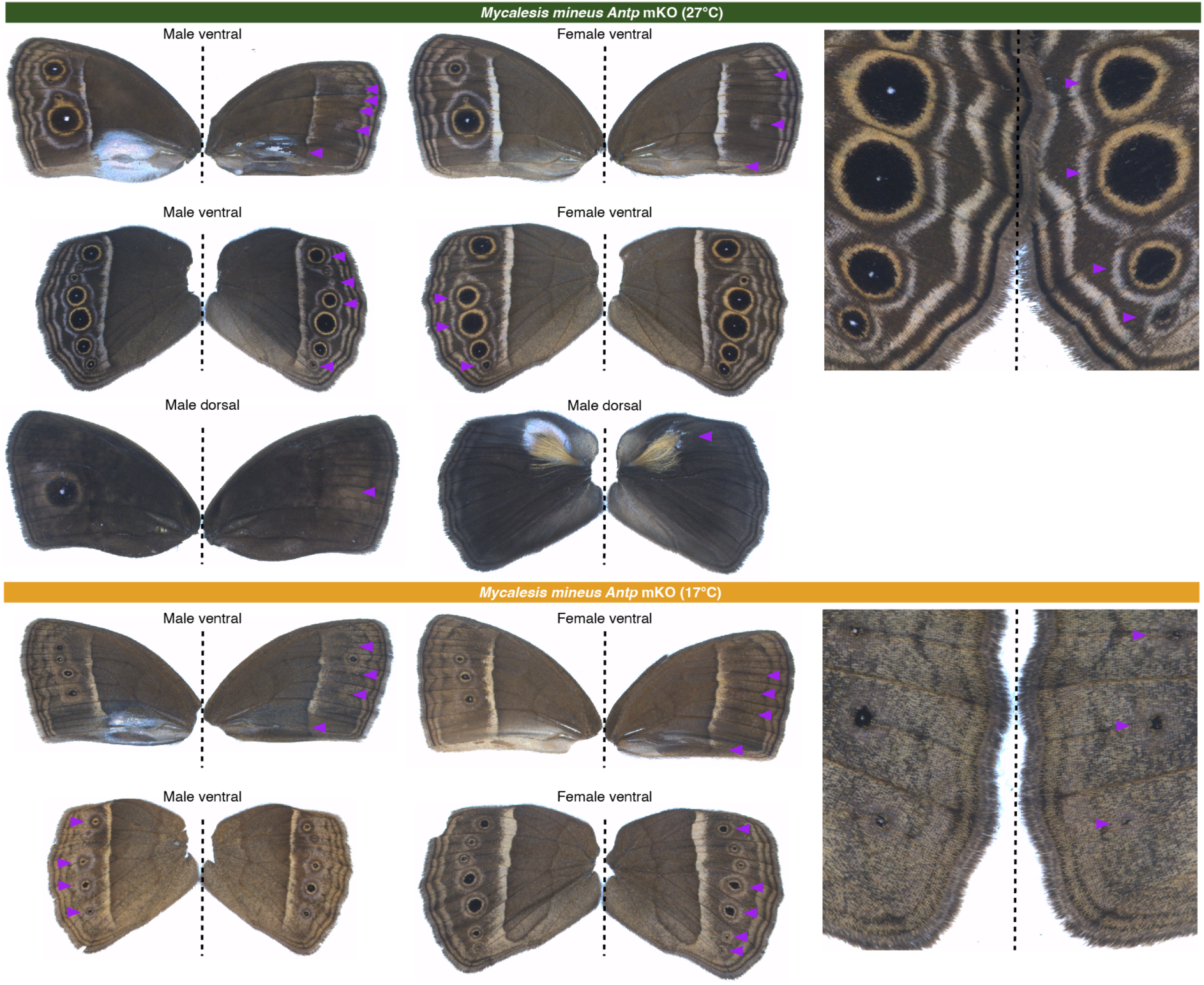
Representative phenotypes of *Antp* mKO crispants in *M. mineus*. A dotted line separates left and right sides of the same individual. Purple arrowheads denote mutant phenotypes.

**fig. S11.**
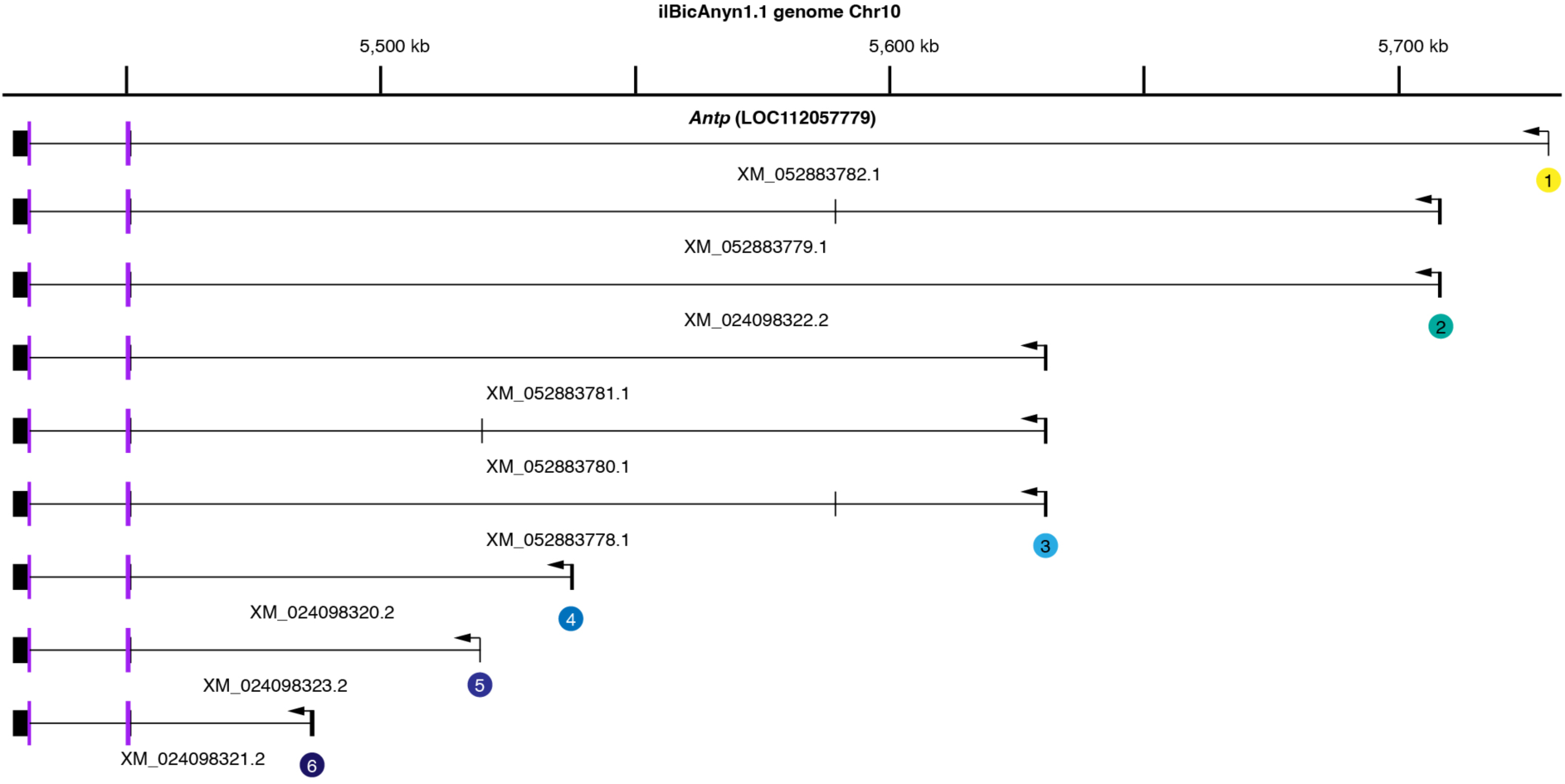
Gene model of *B. anynana Antp*. Gene model is adopted from ilBicAnyn1.1 genome annotation, with NCBI gene and transcript IDs. Six alternative promoters are labeled. CDS is in purple.

**fig. S12.**
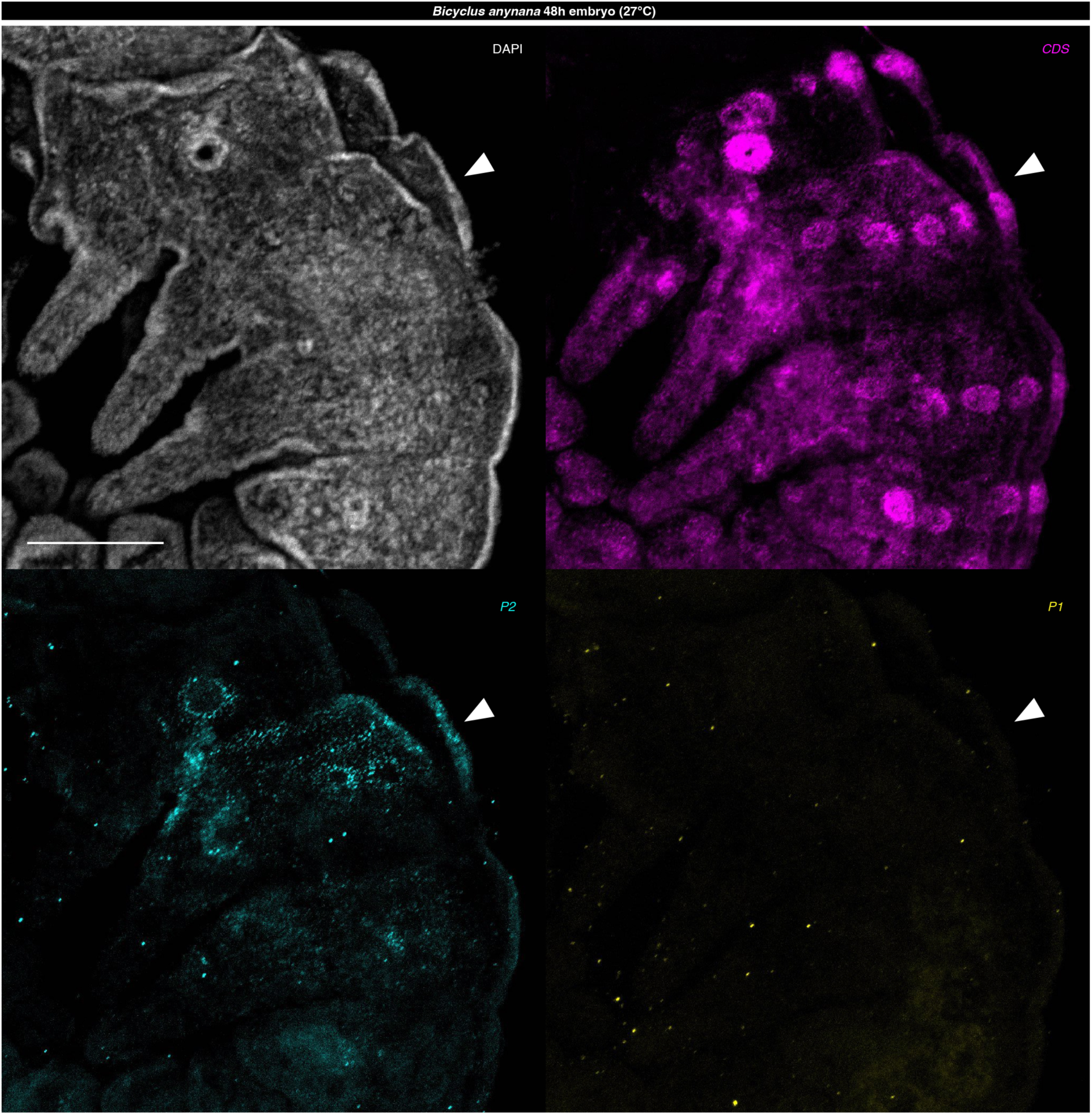
HCR staining of *Antp* features in *B. anynana* 48h embryos. High-magnification images of the HCR co-staining show spatial expression of Antp features in *B. anynana* 48h embryos. Arrowheads denote T2 segments. Scale bar: 100 microns.

**fig. S13.**
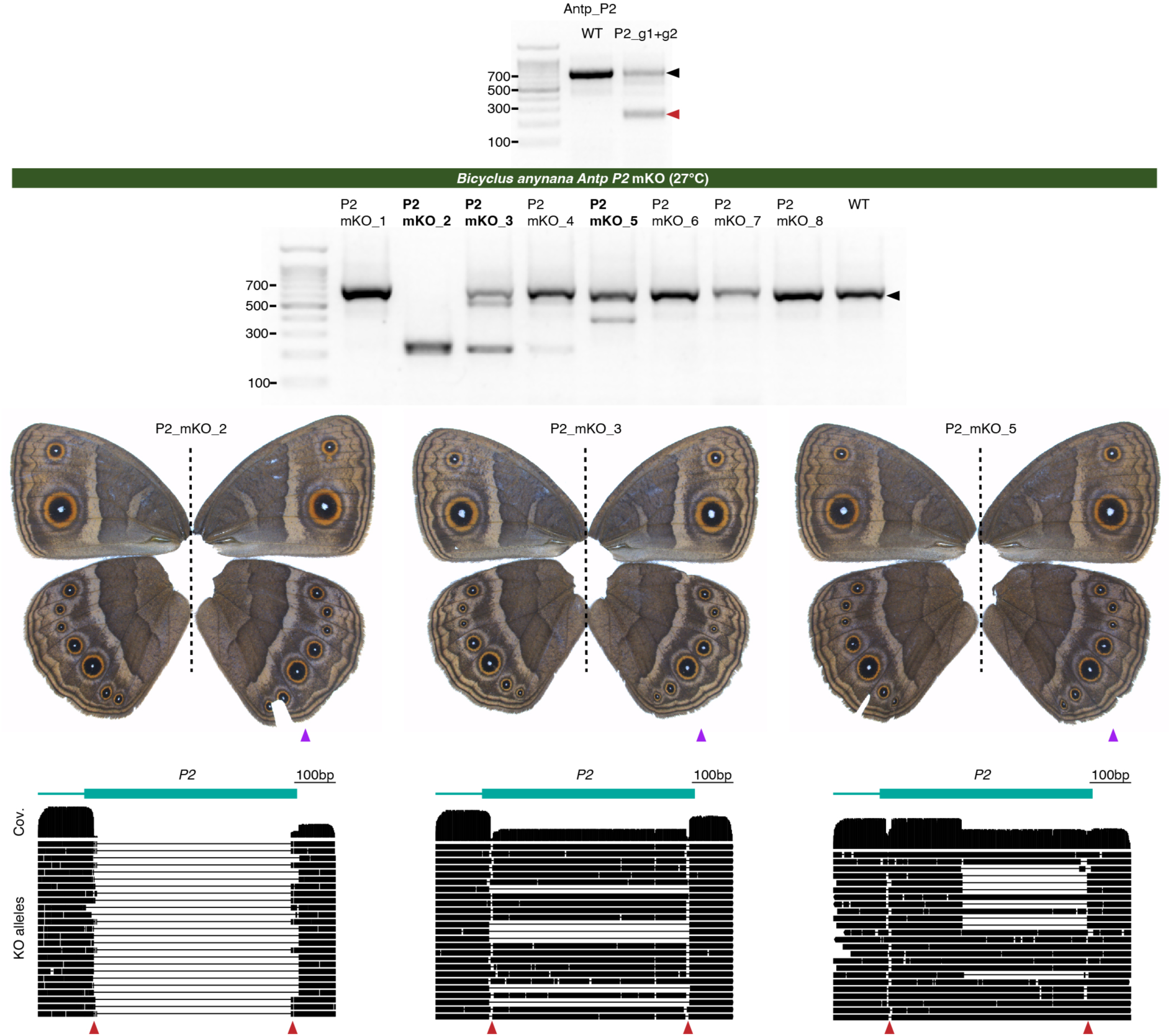
*Antp P2* mKO crispants in *B. anynana*. Two guide RNAs co-injected efficiently induced long deletions around *Antp P2* in vivo, confirmed in pooled injected eggs (top panel). The WT bands and mutant bands are denoted by black and red arrowheads, respectively. Genomic DNA was extracted from right hindwings of eight mKO crispants, and subjected to PCR and gel electrophoresis to detect potential long deletions (mid panel). Three crisptants showing clear long deletions were genotyped via Nanopore amplicon sequencing. A dotted line separates left and right sides of the same individual. Purple arrowhead denote the wings used for genotyping, and red arrowheads denote guide RNA cut sites. Sequence coverage (Cov.) across the amplicon and major KO alleles are shown.

**fig. S14.**
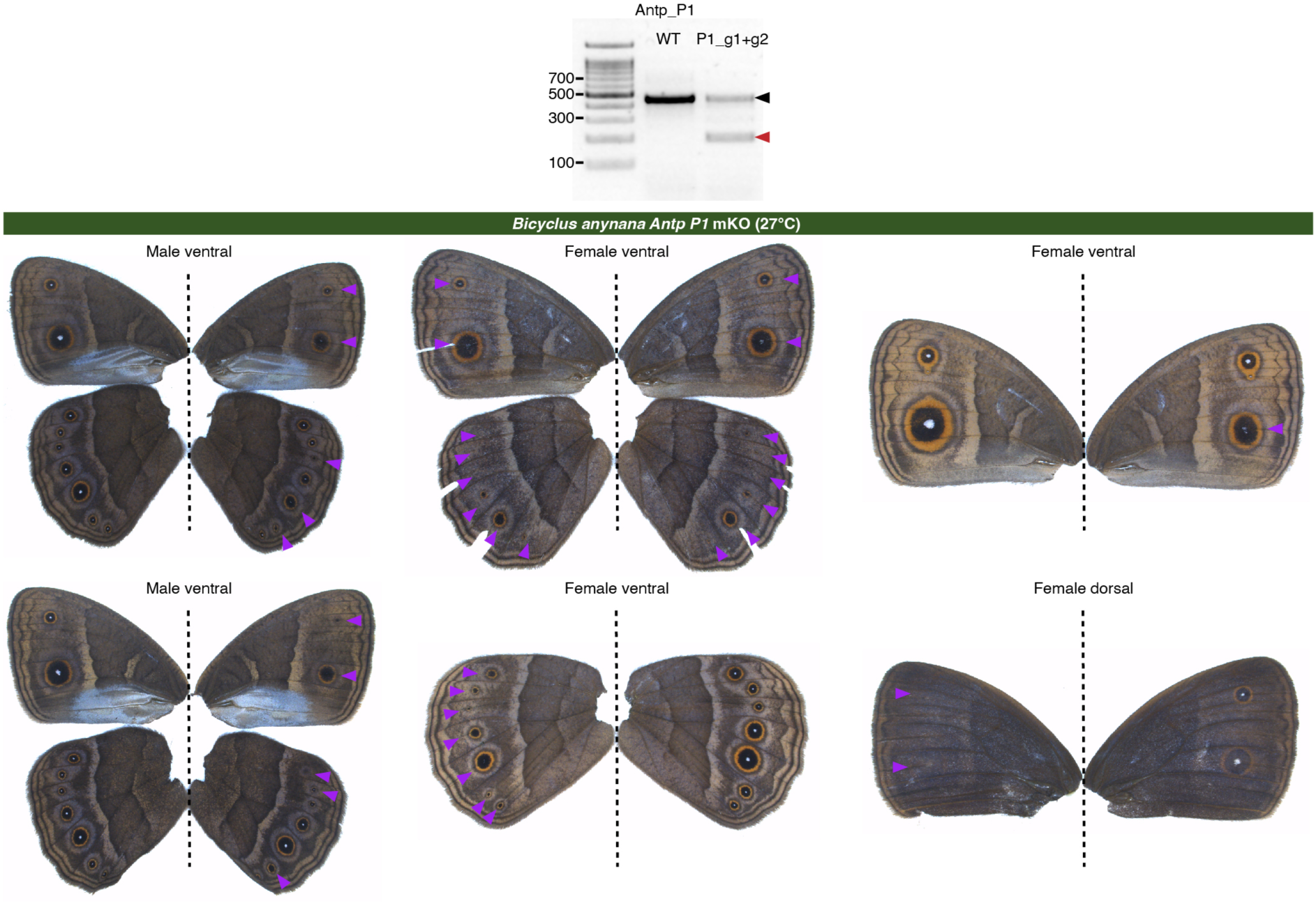
*Antp P1* mKO crispants in *B. anynana*. Two guide RNAs co-injected efficiently induced long deletions around *Antp P1* in vivo, confirmed in pooled injected eggs (top panel). The WT bands and mutant bands are denoted by black and red arrowheads, respectively. A dotted line separates left and right sides of the same individual. Purple arrowheads denote mutant phenotypes.

**fig. S15.**
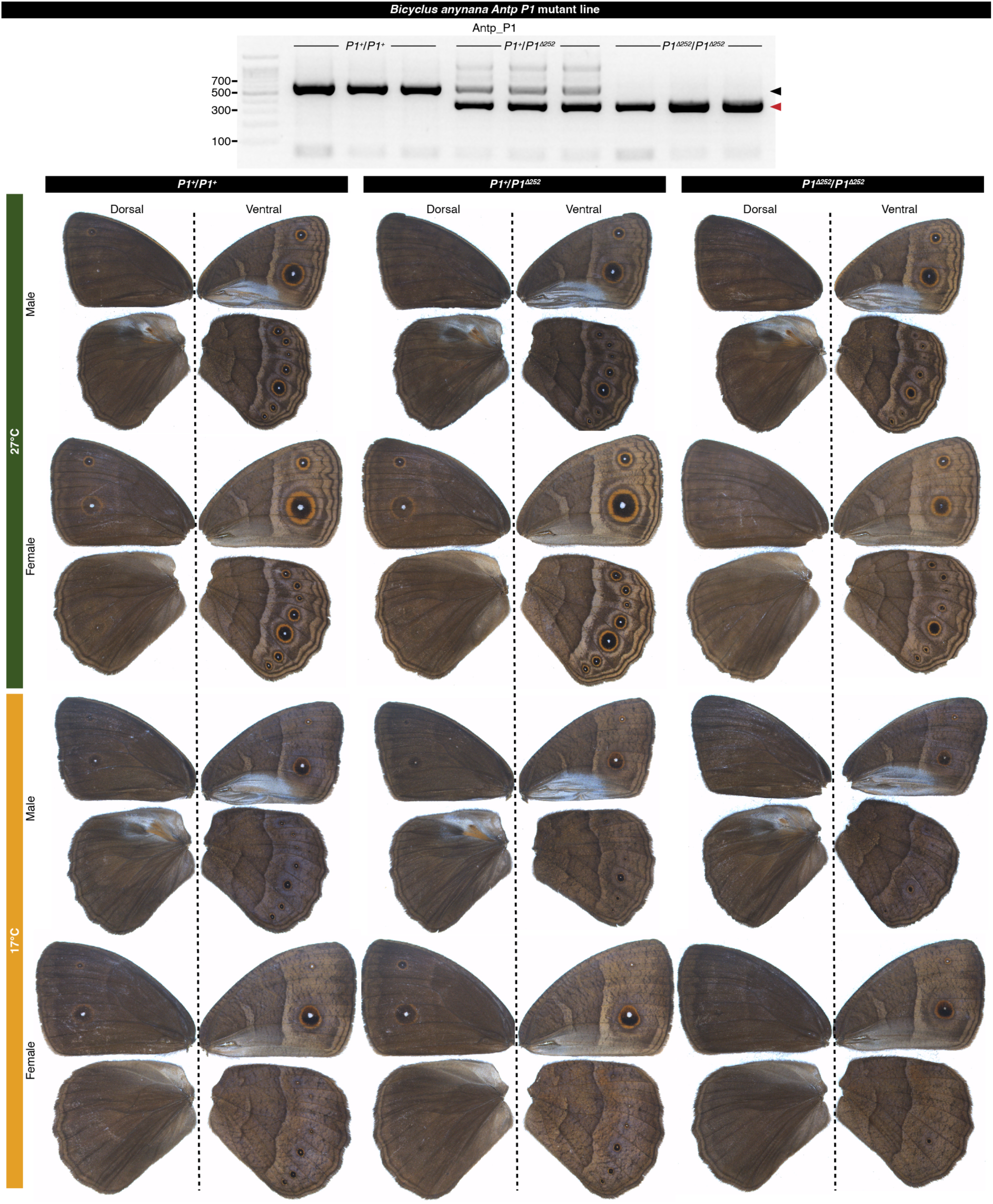
An *Antp P1* mutant line in *B. anynana*. A mutant line was generated carrying a 252bp deletion. The gel image shows clear differentiation of PCR bands across WT, mutant heterozygotes, and mutant homozygotes. The WT bands and mutant bands are denoted by black and red arrowheads, respectively. Representative wings for each genotype in both seasonal forms are shown. A dotted line separates dorsal (left) and ventral (right) sides of the same individual.

**fig. S16.**
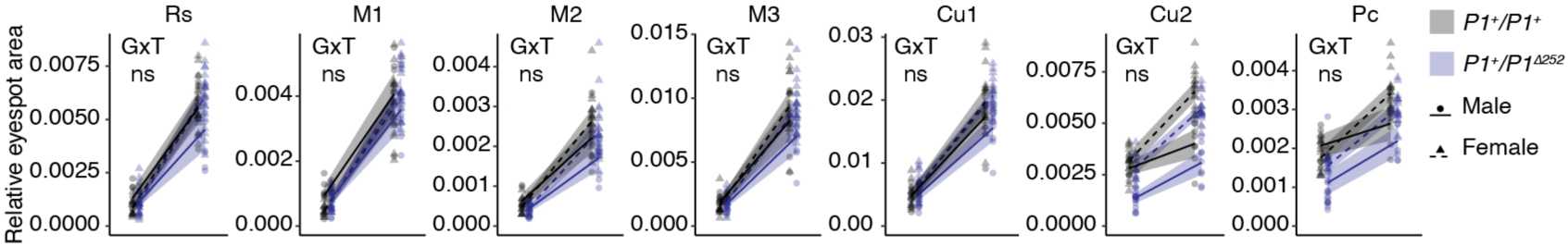
*Antp P1* mutant heterozygotes did not exhibit significant changes in plasticity levels. Changes in eyespot size plasticity levels were assessed across sib-paired WT (*P1^+^*/*P1^+^*) and mutant heterozygotes (*P1^+^*/*P1^Δ252^*) in an ANCOVA, indicated by a significant (p<0.05) genotype (G) x temperature (T) interaction. Shaded line: Mean value with 95CI. ns, not significant; *p<0.05; **p<0.01; ***p<0.001.

### Supplementary Tables

**Table S1.** Summary of the temperature shift experiments.

| Shift scheme<sup>a</sup> | N<sup>b</sup> | Shift start<sup>c</sup> | Shift end<sup>c</sup> | Shift duration<sup>c</sup> |
| --- | --- | --- | --- | --- |
| WS | (35) | / | / | / |
| L5-1 | 21(20) | 0am at 5th instar day0 | 0am at day5 post-shift | 5 day |
| L5-2 | 20(20) | 0am at 5th instar day2 | 0am at day5 post-shift | 5 day |
| L5-3 | 12(12) | 0am at 5th instar day4 | 0am at day5 post-shift | 5 day |
| Wr | 24(24) | 1am | 21.0pm ($\pm 0.6$h) | 44.0h ($\pm 0.6$h) |
| PP | 24(23) | 1.1am ($\pm 0.4$h) | 23.8pm ($\pm 0.4$h) | 46.7h ($\pm 0.4$h) |
| P15 | 22(22) | 21.8pm ($\pm 0.2$h) | 9.8am ($\pm 0.2$h) | 60.0h |
| Wr+PP | 23(23) | 1am | 16.3pm ($\pm 1.0$h) | 87.3h ($\pm 1.0$h) |
| Wr+PP+P15 | 24(23) | 1am | 19.1pm ($\pm 0.7$h) | 6.8 day ($\pm 0.03$ day) |
| L5 | 25(24) | 0am at 5th instar day0 | within 6h post-pupation | 20.4 day ($\pm 0.6$ day) |
| DS | (27) | / | / | / |

<sup>a</sup>Wet season (WS) and and dry season (DS) animals were reared constantly at 27°C and 17°C, respectively.
<sup>b</sup>Number of animals shifted, values in brackets are those survived till adulthood and used for eyespot analysis.
<sup>c</sup>Values in brackets are 95% CI, values without brackets are constants.

**Table S2.** Summary of the differential gene expression analysis.

| Comparisons<sup>a</sup> | DEGs<sup>a</sup> | Systemic plasticity |  |  | Eyespot-specific plasticity |  | Shortlisted eyespot-specific plastic genes |  |  |
| --- | --- | --- | --- | --- | --- | --- | --- | --- | --- |
|  | padj<0.05 | Plastic (padj<0.05) across WSCvsDSC | Homodirectional (systemic) plasticity | % | Not plastic (padj≥0.05) across WSCvsDSC | % | padj<0.01 | llog2FC>0.8 | Mean TPM>2 |
| Wr60_WSEvsDSE | 3460 | 1938 | 1934 | 56% | 1522 | 44% | 868 | 128 | 54 |
| Wr60_WSCvsDSC | 2745 |  |  |  |  |  |  |  |  |
| Wr60_WSEvsWSC | 1983 |  |  |  |  |  |  |  |  |
| Wr60_DSEvsDSC | 2233 |  |  |  |  |  |  |  |  |
| P15_WSEvsDSE | 3519 | 2232 | 2225 | 63% | 1287 | 37% | 683 | 136 | 54 |
| P15_WSCvsDSC | 2892 |  |  |  |  |  |  |  |  |
| P15_WSEvsWSC | 848 |  |  |  |  |  |  |  |  |
| P15_DSEvsDSC | 379 |  |  |  |  |  |  |  |  |

<sup>a</sup>Wr60, 60% wanderer stage; P15, 15% pupal stage; WS, wet season; DS, dry season; E, eyespot; C, control; DEG, differentially expressed gene.

**Table S3.**
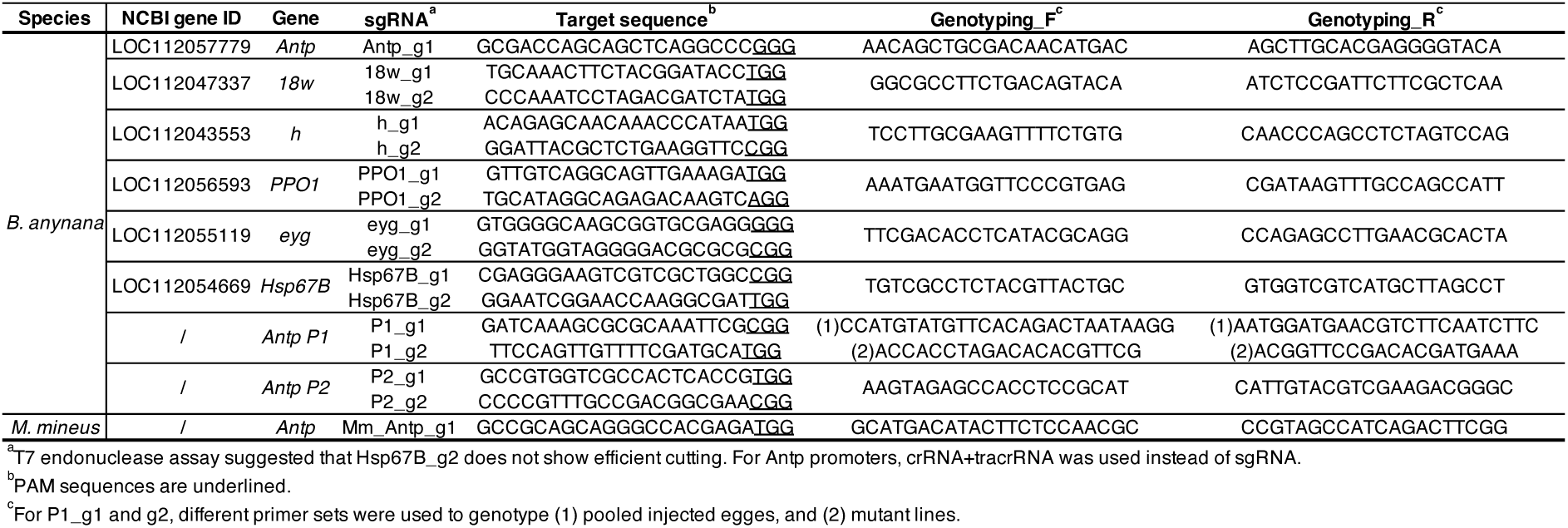
Guide RNA target sequences and genotyping primers.

**Table S4.**
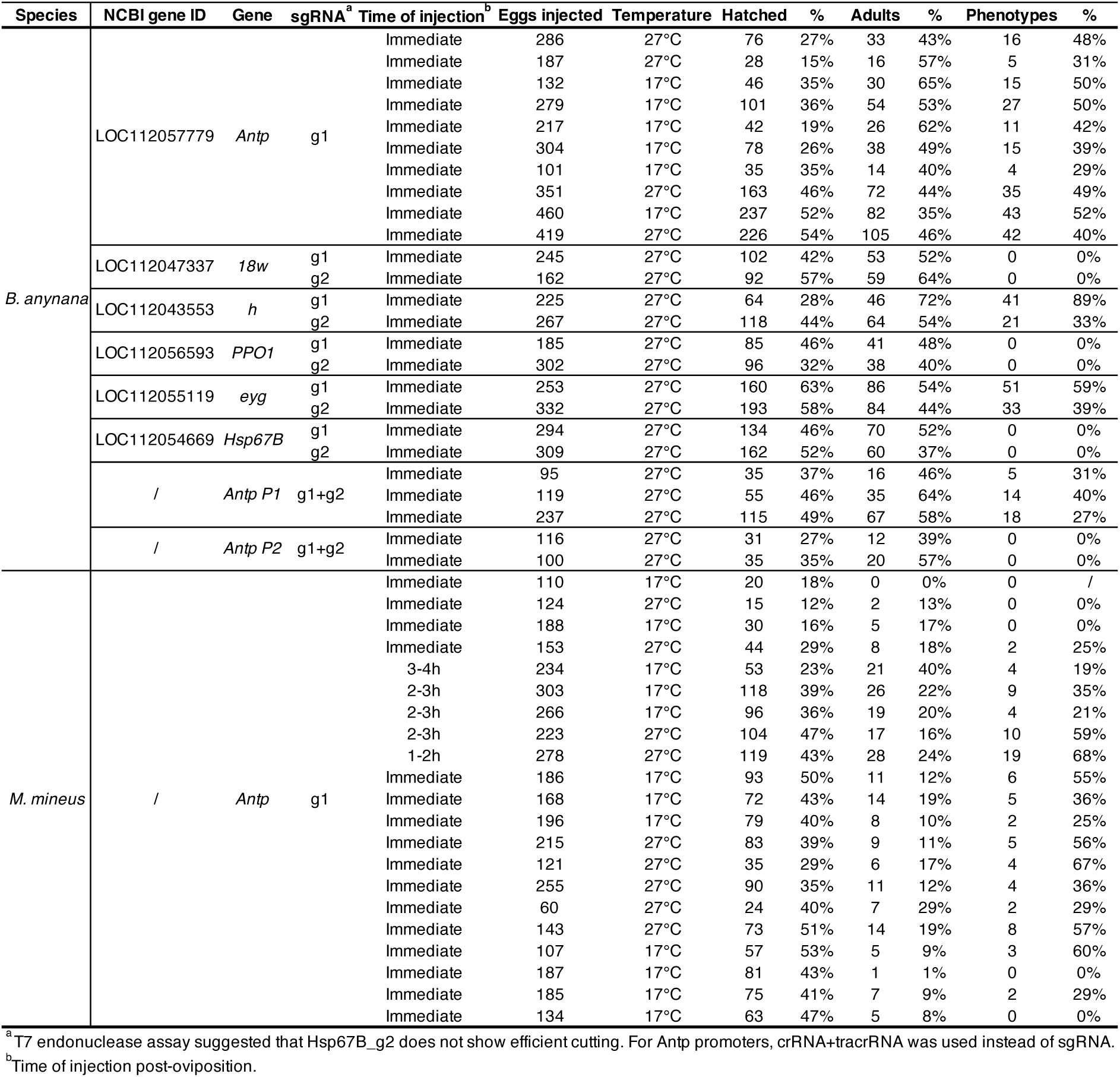
CRISPR injection statistics.

**Table S5.** Sanger genotyping of *eyg* and *h*.

| **Mutant ID** | **Gene** | **Tissue** | **sgRNA** | **Indel rate<sup>b</sup>** |
| --- | --- | --- | --- | --- |
| eyg_g1_12 | *eyg* | LF | eyg_g1 | 74% |
| eyg_g1_13 | *eyg* | LH | eyg_g1 | 94% |
| eyg_g1_14 | *eyg* | RH | eyg_g1 | 73% |
| h_g1_4 | *h* | RF | h_g1 | 88% |
| h_g2_3 | *h* | LH | h_g2 | 65% |
| h_g2_4 | *h* | LH | h_g2 | 72% |

<sup>a</sup>L, left; R, right; F, forewing; H, hindwing.
<sup>b</sup>Calculated from the ICE Synthego tool.

**Table S6.** ANCOVA statistics for the eyespot expression diameter of Antp protein.

| Species | Eyespot | Trans.<sup>a</sup> | Factors |  |  |  |  |  |  |  |
| --- | --- | --- | --- | --- | --- | --- | --- | --- | --- | --- |
|  |  |  | Wing size |  |  |  | Temperature |  |  |  |
|  |  |  | DFn | DFd | F | p | DFn | DFd | F | p |
| *B. anynana* | Rs | / | 1 | 24 | 7.2 | 1.3E-02 | 1 | 24 | 67.9 | 1.9E-08 |
|  | M1 | / | 1 | 24 | 8.0 | 9.0E-03 | 1 | 24 | 156.9 | 5.1E-12 |
|  | M2 | / | 1 | 24 | 15.8 | 5.6E-04 | 1 | 24 | 45.6 | 5.5E-07 |
|  | M3 | / | 1 | 24 | 19.6 | 1.8E-04 | 1 | 24 | 119.4 | 8.4E-11 |
|  | Cu1 | / | 1 | 24 | 32.7 | 6.9E-06 | 1 | 24 | 133.2 | 2.8E-11 |
|  | Cu2 | / | 1 | 24 | 10.9 | 3.0E-03 | 1 | 24 | 39.2 | 1.8E-06 |
|  | Pc | / | 1 | 24 | 15.5 | 6.3E-04 | 1 | 24 | 74.1 | 8.4E-09 |
| *M. mineus* | Rs | / | 1 | 17 | 6.5 | 2.1E-02 | 1 | 17 | 6.4 | 2.1E-02 |
|  | M1 | / | 1 | 17 | 0.3 | 5.8E-01 | 1 | 17 | 0.6 | 4.7E-01 |
|  | M2 | / | 1 | 17 | 0.8 | 4.0E-01 | 1 | 17 | 7.2 | 1.6E-02 |
|  | M3 | / | 1 | 17 | 1.0 | 3.4E-01 | 1 | 17 | 2.1 | 1.6E-01 |
|  | Cu1 | / | 1 | 17 | 9.0 | 8.0E-03 | 1 | 17 | 5.9 | 2.7E-02 |
|  | Cu2 | S | 1 | 17 | 11.0 | 4.0E-03 | 1 | 17 | 24.1 | 1.3E-04 |
|  | Pc | S | 1 | 17 | 3.3 | 8.5E-02 | 1 | 17 | 10.3 | 5.0E-03 |

<sup>a</sup>Data was square root (S) transformed, when necessary, to meet homogeneity of variance criteria, as determined by a Levene's test.

**Table S7.**
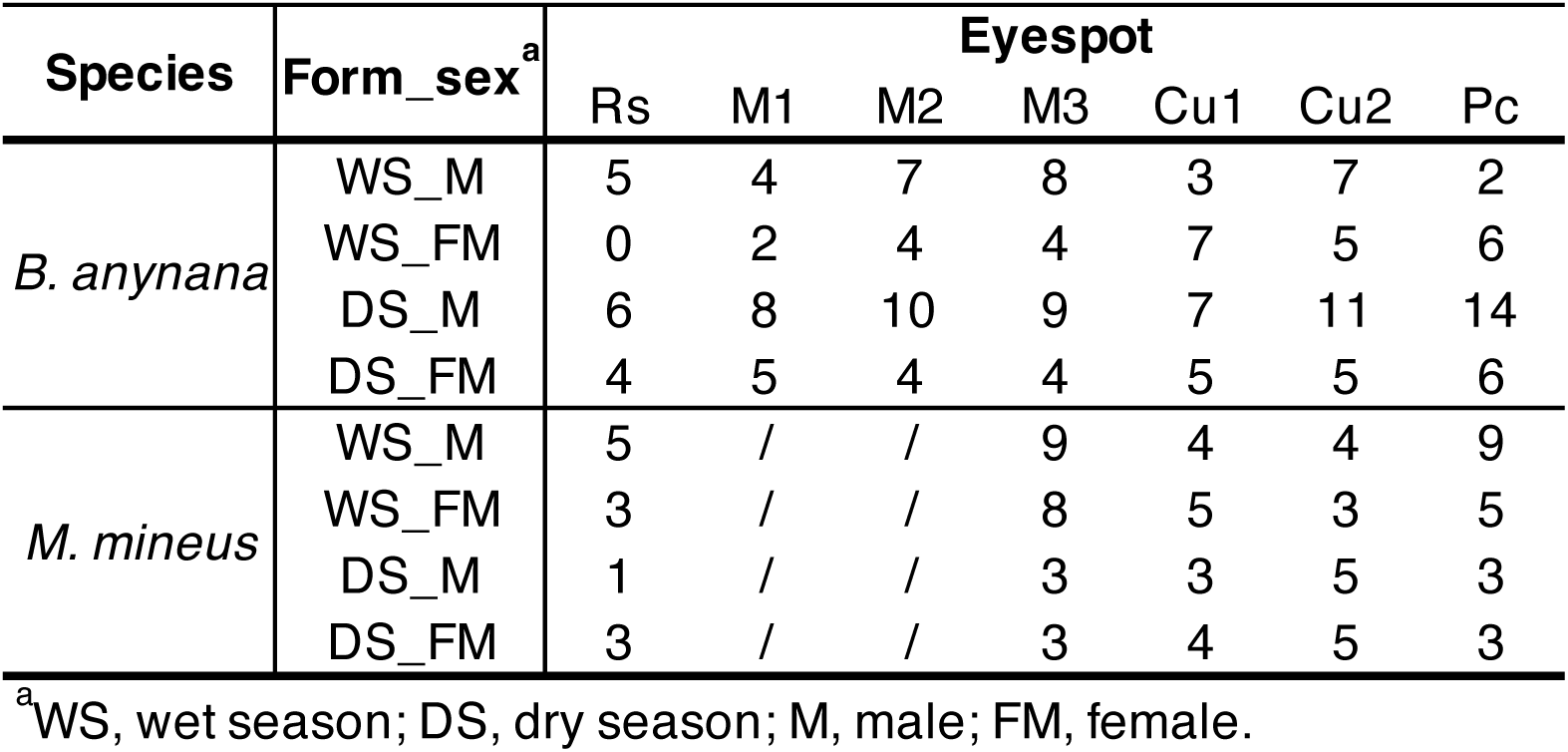
Number of eyespots with paired phenotypes from the *Antp* mKO experiments.

| Species | Form_sex<sup>a</sup> | Eyespot |  |  |  |  |  |  |
| --- | --- | --- | --- | --- | --- | --- | --- | --- |
|  |  | Rs | M1 | M2 | M3 | Cu1 | Cu2 | Pc |
| *B. anynana* | WS_M | 5 | 4 | 7 | 8 | 3 | 7 | 2 |
|  | WS_FM | 0 | 2 | 4 | 4 | 7 | 5 | 6 |
|  | DS_M | 6 | 8 | 10 | 9 | 7 | 11 | 14 |
|  | DS_FM | 4 | 5 | 4 | 4 | 5 | 5 | 6 |
| *M. mineus* | WS_M | 5 | / | / | 9 | 4 | 4 | 9 |
|  | WS_FM | 3 | / | / | 8 | 5 | 3 | 5 |
|  | DS_M | 1 | / | / | 3 | 3 | 5 | 3 |
|  | DS_FM | 3 | / | / | 3 | 4 | 5 | 3 |

<sup>a</sup>WS, wet season; DS, dry season; M, male; FM, female.

**Table S8.**
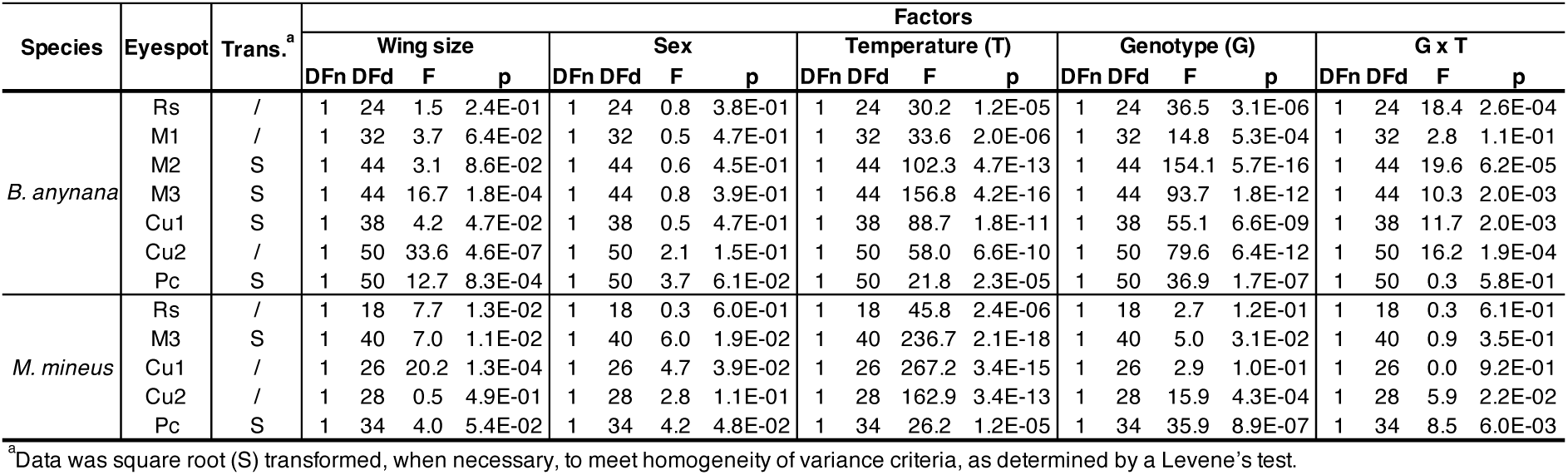
ANCOVA statistics for the *Antp* mKO experiments.

**Table S9.**
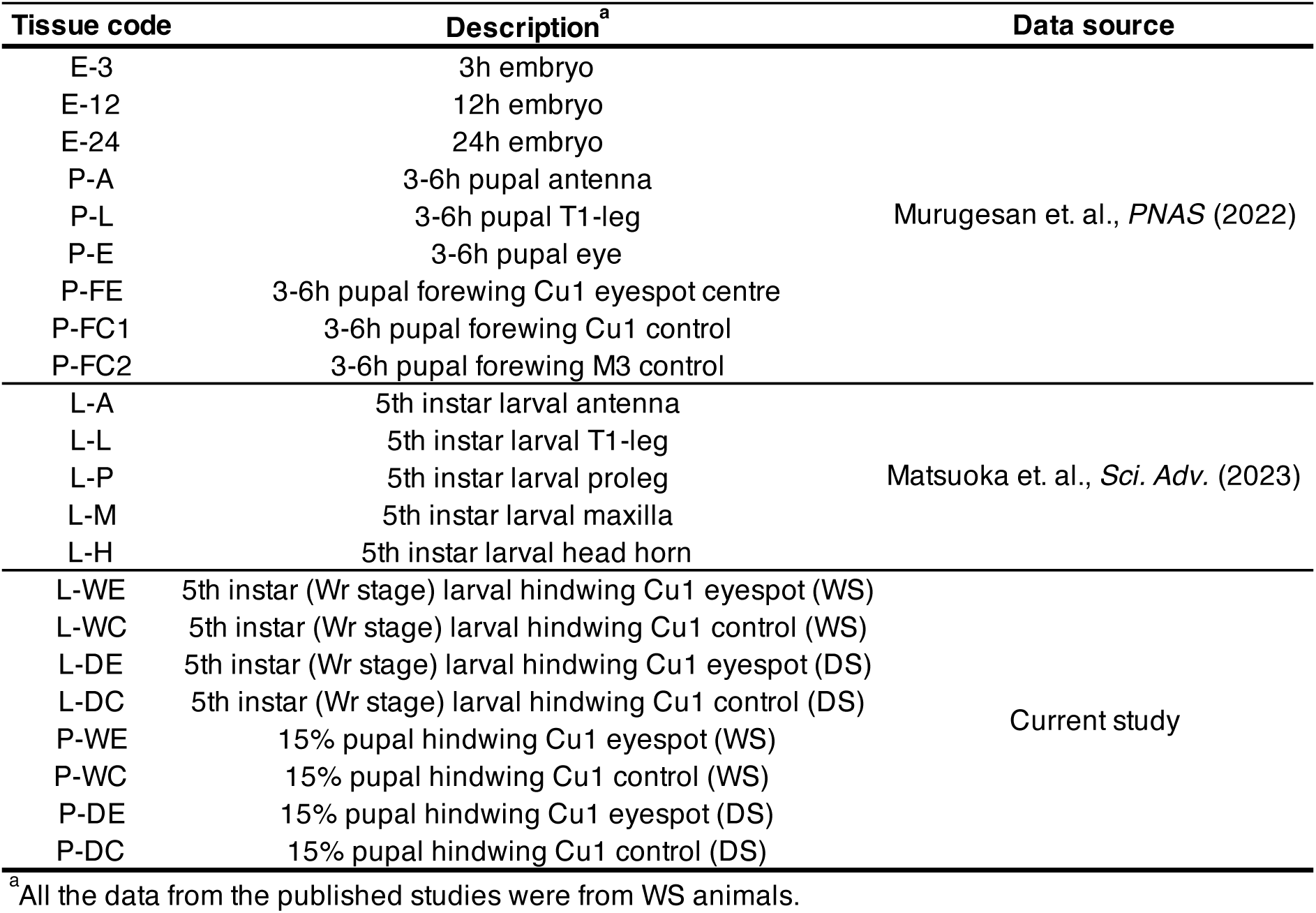
Abbreviation of tissue codes and data sources in the promoter usage analysis.

**Table S10.** Number of eyespots from sib-paired *P1* mutants for eyespot measurements.

| Species | Form <sup>a</sup> | Sex <sup>a</sup> | Genotype | Eyespot |  |  |  |  |  |  |
| --- | --- | --- | --- | --- | --- | --- | --- | --- | --- | --- |
|  |  |  |  | Rs | M1 | M2 | M3 | Cu1 | Cu2 | Pc |
| *B. anynana* | WS | M | *P1<sup>+</sup>/P1<sup>+</sup>* | 9 | 9 | 9 | 9 | 9 | 8 | 7 |
|  |  |  | *P1<sup>+</sup>/P1<sup>Δ252</sup>* | 9 | 9 | 9 | 9 | 9 | 9 | 7 |
|  |  |  | *P1<sup>Δ252</sup>/P1<sup>Δ252</sup>* | 8 | 8 | 8 | 8 | 8 | 8 | 8 |
|  |  | FM | *P1<sup>+</sup>/P1<sup>+</sup>* | 18 | 18 | 18 | 18 | 18 | 18 | 18 |
|  |  |  | *P1<sup>+</sup>/P1<sup>Δ252</sup>* | 21 | 21 | 21 | 21 | 21 | 21 | 18 |
|  |  |  | *P1<sup>Δ252</sup>/P1<sup>Δ252</sup>* | 6 | 6 | 6 | 6 | 6 | 5 | 5 |
|  | DS | M | *P1<sup>+</sup>/P1<sup>+</sup>* | 8 | 8 | 8 | 8 | 8 | 8 | 8 |
|  |  |  | *P1<sup>+</sup>/P1<sup>Δ252</sup>* | 17 | 17 | 17 | 17 | 17 | 17 | 17 |
|  |  |  | *P1<sup>Δ252</sup>/P1<sup>Δ252</sup>* | 8 | 8 | 8 | 8 | 8 | 7 | 6 |
|  |  | FM | *P1<sup>+</sup>/P1<sup>+</sup>* | 12 | 12 | 12 | 12 | 12 | 12 | 12 |
|  |  |  | *P1<sup>+</sup>/P1<sup>Δ252</sup>* | 16 | 16 | 16 | 16 | 16 | 16 | 16 |
|  |  |  | *P1<sup>Δ252</sup>/P1<sup>Δ252</sup>* | 14 | 14 | 14 | 14 | 14 | 14 | 14 |

<sup>a</sup>WS, wet season; DS, dry season
<sup>b</sup>M, male; FM, female.

**Table S11.**
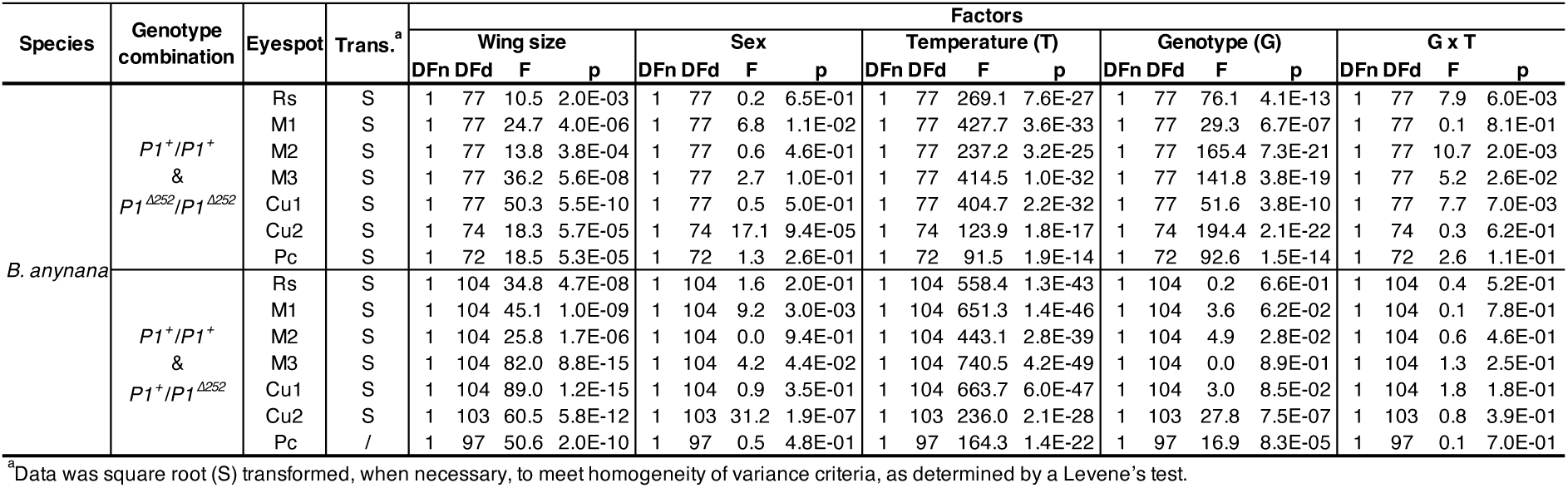
ANCOVA statistics for the *P1* mutant line.

